# Recommendations for Quantitative Data-Independent Acquisition (DIA) Proteomics using Controlled Quantitative Experiments (CQEs)

**DOI:** 10.1101/2025.09.22.677725

**Authors:** Andrew M Frey, Frances Sidgwick, Andrew Porter, Pawel Palmowski, Matthias Trost

## Abstract

Advances in mass spectrometry and associated computational tools have resulted in the wide-spread adoption of Data-Independent Acquisition (DIA) for proteomics. Experiments using state-of-the-art instrumentation and specialised modes of acquisition report ever increasing throughput and protein identification rates. However, experiments to test the efficacy of protein quantitation in these novel methods is surprisingly rare -especially in novel methods where low sample input or high-throughput is considered. Here, we performed a series of controlled quantitative experiments (CQEs) on two commonly used mass spectrometry platforms (Exploris480 and timsTOF HT), using defined mixtures of human, yeast, and bacterial proteomes. Peptide spectrum matching and initial quantification was performed using DIA-NN, with a series of post-processing options also compared. Overall, we identified >10,000 unique proteins and >160,000 precursors across samples with excellent accuracy and precision. Additional post-processing was found to have drastic impacts on ID rates, quantitative precision and accuracy, and false quantitative rate (FQR). These impacts were particularly notable in experiments using low amounts of sample. In optimal loading conditions, data processing could reduce FQR from ∼0.1 to 0 % with minimal losses in ID rates. Ultimately, we present a series of recommendations for proteomics data processing, and advice on the use of CQEs and resulting metrics when assessing quantitative efficacy in novel modes of acquisition.

## Introduction

Quantitative proteomics has made substantial progress over the past decade, with mass spectrometry (MS) becoming a cornerstone of proteomic research (1). This development has cemented MS’s role as an essential tool in biological and biomedical studies. Despite these advancements, achieving efficacious quantitative analysis across the vast dynamic range of the proteome remains a major challenge. The proteome spans an immense range: the cellular proteome covers seven orders of magnitude, while plasma proteomes extend over ten (2, 3). Progress in the field has been propelled by innovations in hardware and data acquisition technologies, pushing proteomics towards more sophisticated applications, particularly in clinical settings for disease diagnostics and biomarker discovery.

In response to these demands, there has been a shift in proteomics from focusing on detecting as many peptides and proteins as possible to prioritising throughput and quantification. Additionally, a transition from data-dependent acquisition (DDA) to data-independent acquisition (DIA) strategies has become more prevalent. DIA LC-MS/MS is now the cornerstone MS acquisition method utilised in proteomic analysis, with recent advancements in software tools and faster mass spectrometers driving its expansion (4). Innovations in MS instrumentation, particularly in the sensitivity and speed of mass analysers, and the adoption of shorter liquid chromatography gradients have contributed to the success of this approach. These combined with enhanced data analysis techniques have led to a preference for label-free quantification (LFQ) methods. LFQ offers multiple benefits, including reduced costs, fewer chemical requirements, and no sample number limitations (5). As the field moves towards high-throughput analysis, the adoption of shorter LC gradients has become more common to reduce overall acquisition times. Shorter LC gradients lead to reduced peak widths, and the speed of the mass analyser becomes crucial for generating enough data points per peak to ensure quantitative accuracy. Although longer LC gradients provide deeper proteome coverage and greater peptide identification, they come at the cost of increased analysis time.

Despite the (now routine) detection of thousands of proteins in recent studies, quantifying low-abundance proteins remains a challenge. Variations in false discovery rate (FDR) thresholds and quantifiable peak identification across studies complicate comparisons, emphasizing the need for rigorous benchmarking of technologies. Managing the FDR is a major statistical challenge in shotgun proteomics. The most common method for addressing this is target-decoy competition (TDC) at the peptide-spectrum match (PSM) level, where observed spectra are searched against a database containing both genuine (target) peptides and decoy peptides. At the peptide level, the top-scoring PSM for each peptide is selected, and FDR is estimated based on this information. In proteomics, an FDR threshold of 1% is commonly accepted, meaning that 1% of the quantifiable proteins will be false positives.

Reporting the total number of proteins identified is a standard output in proteomics experiments. However, relying solely on this metric can be misleading, as it may inflate results and fail to account for reproducibility within datasets. More proteins are typically identified than those that are quantifiable, and the total protein count does not reflect the consistency across replicates. Therefore, it is crucial to assess not only the number of identified proteins but also their quantifiability.

The number of data points collected across each chromatographic peak is usually considered a critical factor for ensuring reproducibility and accurate quantification. It has been reported that at least six data points are typically required to accurately define the peak shape and estimate the peak apex (6). Fewer data points can result in missing values, as well as reduced reproducibility and poor quantitation across replicates. However, the number of data points and protein identifications alone does not indicate the variability and reproducibility within a dataset. To address this, CV values are used to assess reproducibility. The CV expresses variance as a percentage of the mean, with higher values indicating greater dispersion around the mean. Lower CV values suggest good reproducibility, which is vital for ensuring the reliability of the data.

LFQ mixed proteome experiments were initially designed by *Navarro et al* in 2016 (7) and are now utilised by (4, 8, 9). These involve the analysis of multiple proteomes in a single experimental setup without the use of isotopic or chemical labels. Cell lysates from multiple organisms or conditions are first digested and peptides are then pooled together in a single sample, with the ratio of each proteome in the mixture being carefully controlled. Although the absolute concentration of individual proteins is not known, this experimental design allows quantification of different protein ratios for each organism and enables in-depth evaluation of precision and accuracy of the measurements, as well as the proportion of proteins in opposition to the ground truth.

In this manuscript, we describe a series of these controlled quantitative experiments (CQEs), including two- and three-proteome experiments. In addition to describing overall fold change distribution and variance, we calculate additional metrics and use these to interrogate a number of key parameters in MS-proteomics workflows. In addition to MS acquisition, post-processing decisions such as the number of peptides identified per protein, the precursor intensity and data points per peak, and the coefficient of variation (CV) across replicates, to consider when filtering the acquired DIA data to ensure reliable quantitative results, but also while designing and evaluating LC-MS methods. Ultimately, we provide a series of recommendations for optimal data processing and best practice for assurance of quantitative efficacy in DIA-LFQ proteomics.

## Materials and Methods

### Species, Strains, and Culture Methods

HeLa cells were routinely cultured in Dulbecco’s Modified Eagle Medium (DMEM) with 10 % (v/v) Foetal Calf Serum, and 15 mM L-Glutamine, and were passaged at ∼80 % confluency. Prior to collection, cells were washed three times with ice cold PBS and detached by scraping. Cells were centrifuged (5 minutes, 1000 ×g, 4°C) and stored at -80°C. *Escherichia coli* (DH5α) were cultured overnight in LB medium at 37°C in an orbital shaker at 50 rpm. *Saccharomyces cerevisiae* was cultured overnight in Yeast-Peptone-Dextrose (YPD) broth at 37°C in an orbital shaker at 50 rpm. Microbial liquid cultures were collected by centrifugation (10 minutes, 4000 ×g, 4°C), and washed three times with ice cold PBS.

### S-Trap protein digestion

Samples were lysed in 500 µL 5% SDS, 50 mM triethyl ammonium bicarbonate (TEAB). Following protein quantification (Pierce BCA assay, ThermoFisher Scientific) 1 mg of each sample was reduced by adding Tris(2-carboxyethyl)phosphine (TCEP) to a final concentration of 10 mM for 30 min at room temperature followed by alkylation with 20 mM iodoacetamide for 30 min in the dark. Samples were then subjected to suspension trapping and digestion with a commercially available kit (Protifi), according to manufacturer’s instructions. Briefly, samples were acidified by addition of 50 µL of 12% phosphoric acid and diluted with 3.3 mL of S-trap binding buffer: 90% MeOH, 100 mM TEAB, pH 7.1. The acidified samples were then loaded onto the S-trap midi spin column and centrifuged for 1 minute at 4000 ×g. Columns were washed five times with S-trap binding buffer before addition of porcine trypsin (1:10) (Worthington) in 350 µL of 50 mM TEAB to the column. Samples were incubated at 37°C over-night. Peptides were eluted by washing the column with firstly 50 mM TEAB, pH 8.0 (500 µL), followed by 500 µL of 0.2% formic acid (FA) and finally 500 µL of 0.2% FA, 50% acetonitrile. Peptides were then dried under vacuum. Peptides from separate species were resuspended and combined in defined ratios as described in the materials and methods.

### E480 LC-MS methods

Protein digests were dissolved in 0.2% trifluoroacetic acid and injected with an UltiMate 3000 RSLCnano System (Thermo Fisher). Samples were trapped on an Acclaim PepMap 100 C18 LC trap column (Inner diameter: 100 μm × 20 mm, 3 μm, 100 Å) and then separated on an 75cm EASY-Spray nanoLC C18 column (Inner diameter: 75 μm × 500 mm, 2 μm, 100 Å; Thermo-Fisher) at a flow rate of 150 nL/min. Solvent A was water containing 0.1% formic acid, and solvent B was 80% acetonitrile containing 0.1% formic acid. The gradient used for analysis of samples was as follows: Alosolvent B was maintained at 3% for 5 min, followed by an increase from 3% to 35% B over 120 min, 35 to 90% B over 0.5 min, maintained at 90% B for 5 min, followed by a decrease to 3% over 0.5 min and equilibration at 3% for 25 min.

The Exploris 480 mass spectrometer was operated in positive ion, DIA mode. Full MS scan spectra were acquired in the m/z range of 350-1050 with normalised automatic gain control target of 1000%, and maximum injection time set to dynamic, at a resolution of 60,000. Targeted MS2 scan spectra were acquired using either 20, 76 or 152 isolation windows (**Supplemental table S1**). CE set to normalised if not stated. AGC target of 3000%, at a resolution of 30,000 with dynamic injection time. Electrospray voltage was static, and capillary temperature was 275°C, with expected LC peak width of 30s. No sheath and auxiliary gas flow were used

### timsTOF-HT LC-TIMS-MS/MS methods

LC was performed using an Evosep One system with a 15 cm Aurora Elite C18 column with integrated captive spray emitter (IonOpticks), at 50°C. Buffer A was 0.1% formic acid in HPLC water, buffer B was 0.1% formic acid in acetonitrile. Immediately prior to LC-MS, peptides were resuspended in buffer A and a volume equivalent to 500 ng was loaded onto the LC system-specific C18 EvoTips, according to manufacturer instructions, and subjected to the predefined Whisper-Zoom 20 SPD protocol (where the gradient is 0-35 % buffer B, 200 nl/min, for 58 minutes, 20 samples per day were permitted). For high-sensitivity experiments the Whisper-Zoom 40 SPD protocol (40 samples per day permitted) was used, with 250 pg of protein digest loaded. The Evosep One was used in line with a timsTOF-HT mass spectrometer (Bruker) operated in diaPASEF mode. Mass and IM ranges were 300-1200 *m/z* and 0.6-1.45 1/*K_0_*. diaPASEF was performed using 16 variable width IM-*m/z* windows each with two frames, without overlap (**Supplemental table 2)**. TIMS ramp and accumulation times were varied as indicated in the results, though for the most frequently used 66 ms, total cycle time was ∼1.2 seconds. For high sensitivity, a 4-window scheme (**Supplemental table 3**) with 300 ms accumulation time was used. Collision energy was applied in a linear fashion, where ion mobility = 0.6-1.6 1/K_0_, and collision energy = 20 - 59 eV.

### Data Analysis

Raw files were searched using DIA-NN version 1.9.2 (reference DIANN: P31768060) (10) against a Uniprot *H. sapiens, E. coli, and S. cerevisiae* reference database to identify protein groups (containing 30826 proteins-see figure 3, downloaded July 2024) as well as a common contaminants database (11), at 1% false discovery rate with default settings (with the exception that searches for the timsTOF-HT were set to precursors with *m/z* 300-1200). *In silico* digest for spectral library generation was used for library-free search. For high sensitivity (250pg) experiments on the timsTOF-HT, an empirical DIA-spectral library was used. This was generated by subjecting 50 ng of matched samples (equivalent to “bulk” preparations in single-cell experiments) to the same LC-TIMS-MS high sensitivity protocol, analysed by DIA-NN as described directly above. All resulting datasets underwent post-processing as follows: The report.parquet DIA-NN output file was subjected to processing in Rstudio. Key were diannR, tidyverse, ggplot2, plotly, and dyplyr packages. A variety of post-processing options were tested as described in the results section. Common to all analyses was the use of Limma for statistics (12), automated exclusion of contaminants, and only including proteins quantified in at least half (3/5 or 2/3) of replicates for both conditions in a given comparison.

**Figure 1.**
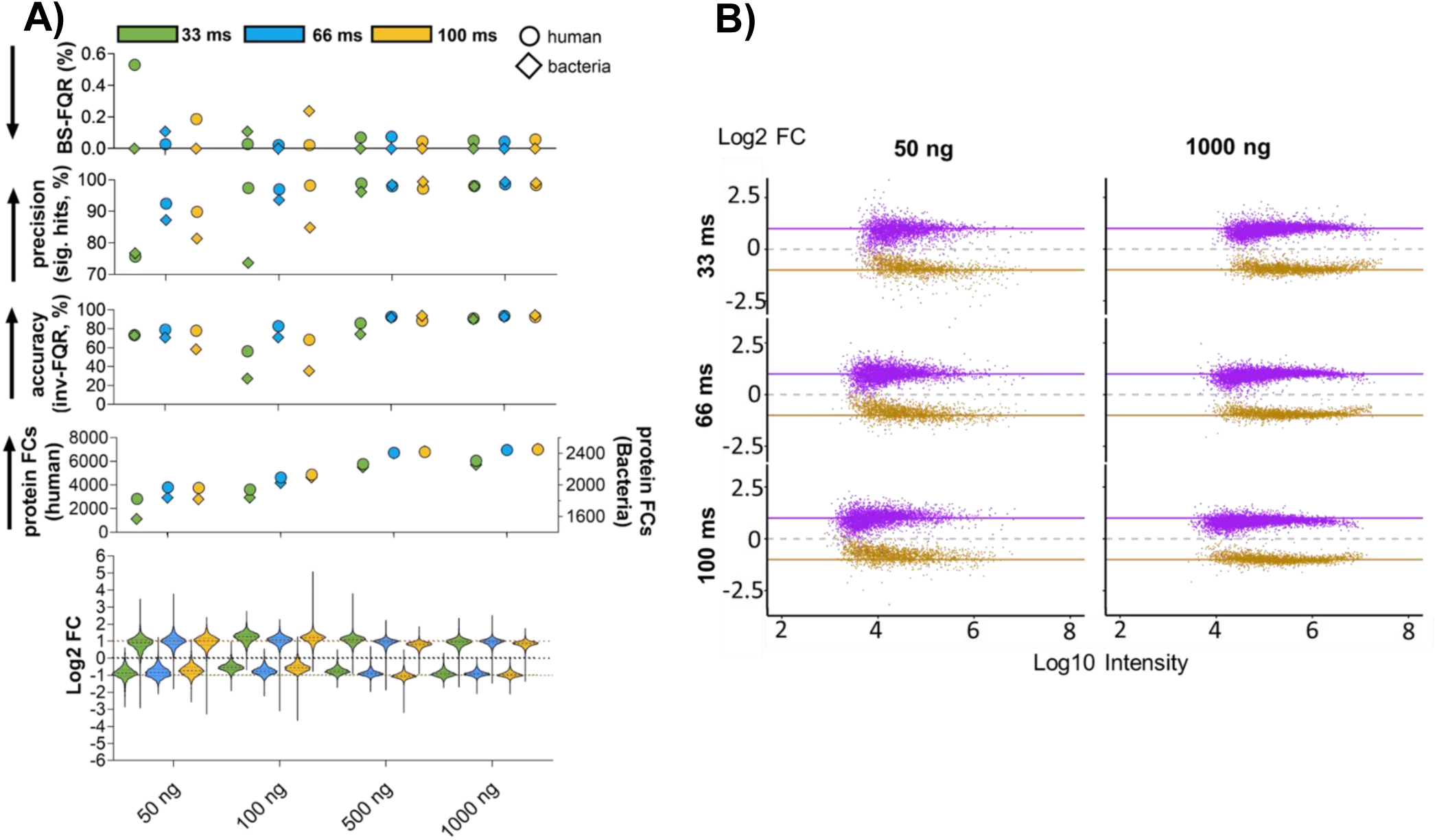
Initial assessment of quantitative efficacy for diaPASEF with variable TART and load. Left: Summary of metrics for TART and load conditions, lower to upper figures; Log2 FC (median, 25 % quartiles, outliers shown, expected FC and zero are indicated by dashed lines), protein group FCs, Significant hits, FQR, and BS-FQR are shown. Arrows indicate the preferable result vector for each metric, i.e. If a higher result is better (arrow pointing up) or worse (arrow pointing down). Right: Distribution of (log10) protein group intensities against (log2) fold change for all TART, highest and lowest load (50 and 1000 ng). Purple = Human, Khaki = Bacteria. Data shown are derived from three replicates.

**Figure 2.**
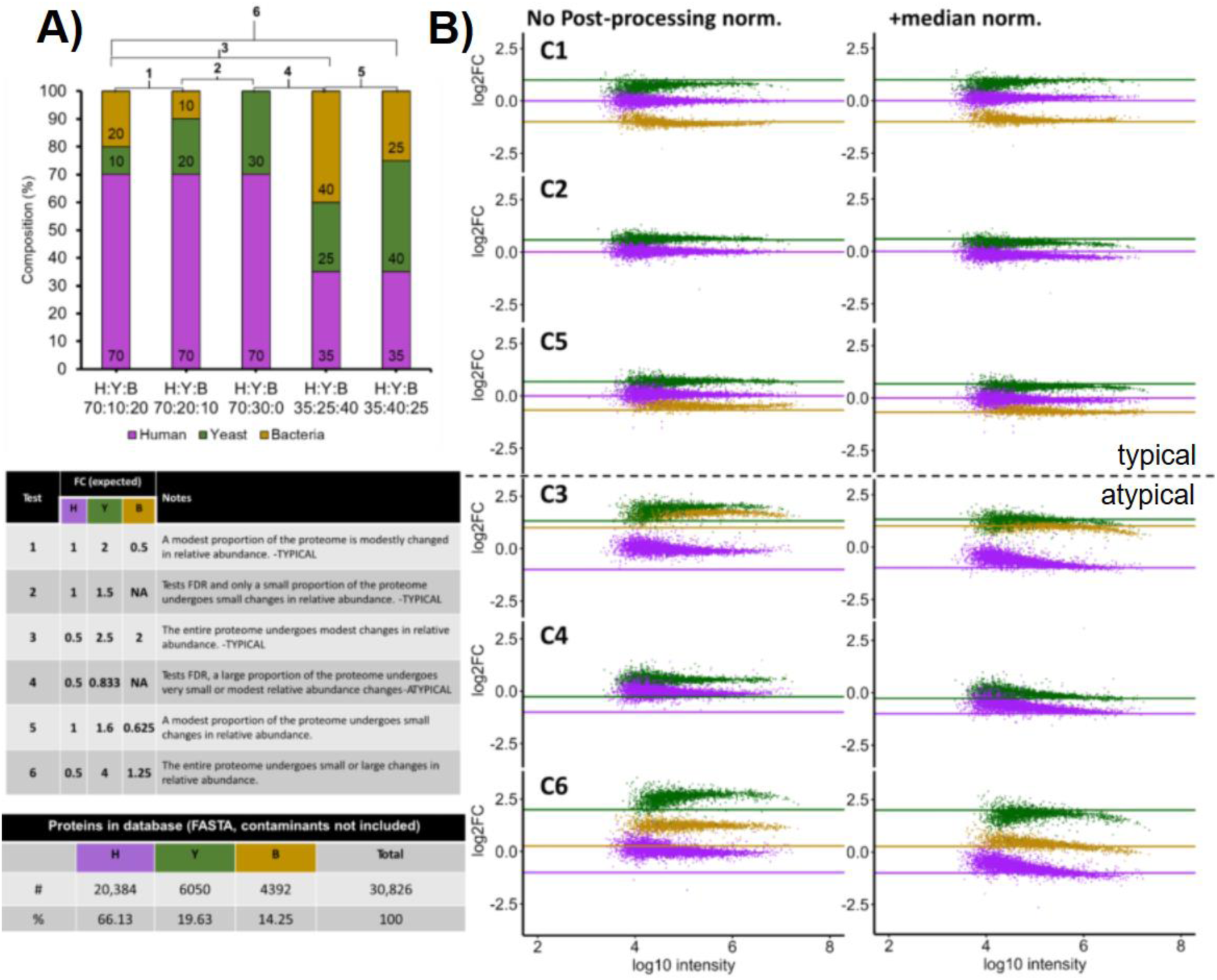
Three species CQEs. A) Rationale for comparisons. B) Median normalisation alters distribution of measured protein group FC (points) around the ground truth (lines). FC shown here is the default post-processing pipeline (2 peptides per protein, protein present in 3/5 replicates in both conditions, no other filtering).

**Figure 3.**
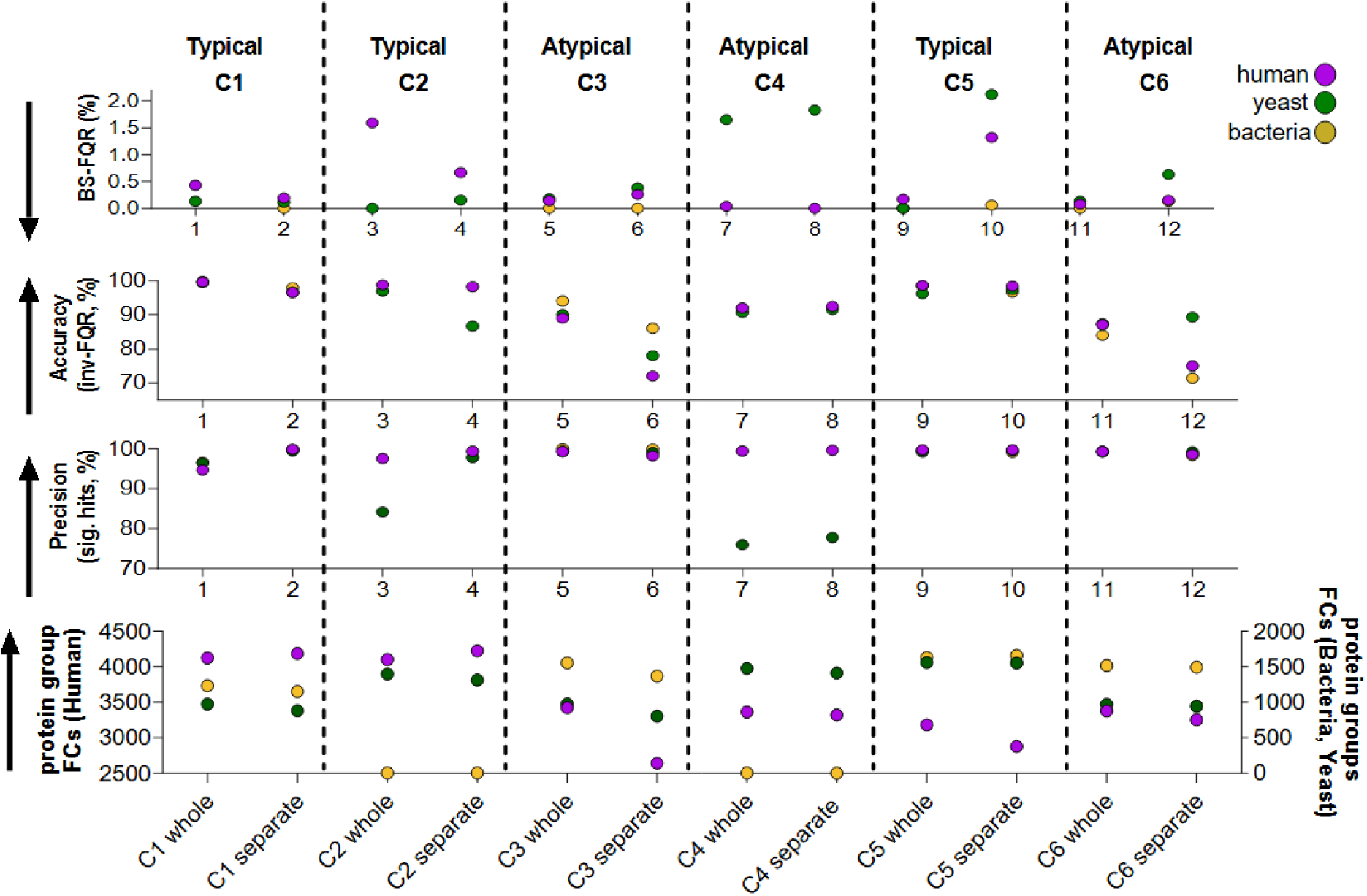
Search space has an impact on quantitative metrics. Raw files were analysed by DIA-NN separately for each condition (separate) or all at the same time (whole). Data shown underwent median normalisation, protein present in 3/5 replicates in both conditions, with protein group CV filtering. Data are derived from 5 replicates.

### Data Availability

The mass spectrometry proteomics data have been deposited to the ProteomeXchange Consortium via the PRIDE (7) partner repository with the dataset identifier PXD068065 for the E480, and PXD068192 for the timsTOF-HT. Reviewers can access the datasets using the following details: For the E480-Project accession; PXD068065, Token: pCaINJrFFruY, for the timsTOF-HT-Project accession: PXD068192, Token: FjZNJmHZyVJV.

## Results

### Optimizing DIA Parameters with Two-Species Controlled Quantitative Experiments (CQEs)

Controlled quantitative experiments (CQEs) provide a robust framework for evaluating both mass spectrometry acquisition parameters and subsequent data analysis workflows. To optimize our data-independent acquisition (DIA) strategy, we employed a two-species CQE using a tryptic digest mixture of human (HeLa) and bacterial (*E. coli*) proteins. The mixtures were prepared at two different ratios of human to bacterial proteins (1:2 and 2:1) while maintaining a constant total protein load.

The performance of the tested acquisition parameters was evaluated using metrics summarised in **Table 1**. Accuracy was quantified as the inverse false quantitative rate (inv-FQR, %) and defined as the proportion of proteins with fold changes within ±25 % of the expected value, a stricter threshold than the ±50 % commonly used in proteomics studies. Precision was assessed as the proportion of statistically significant hits. To address the risk of biologically misleading results, we additionally calculated the biologically significant false quantitative rate (BS-FQR, %), defined as the proportion of statistically significant proteins with fold changes in the incorrect direction.

**Table 1.**
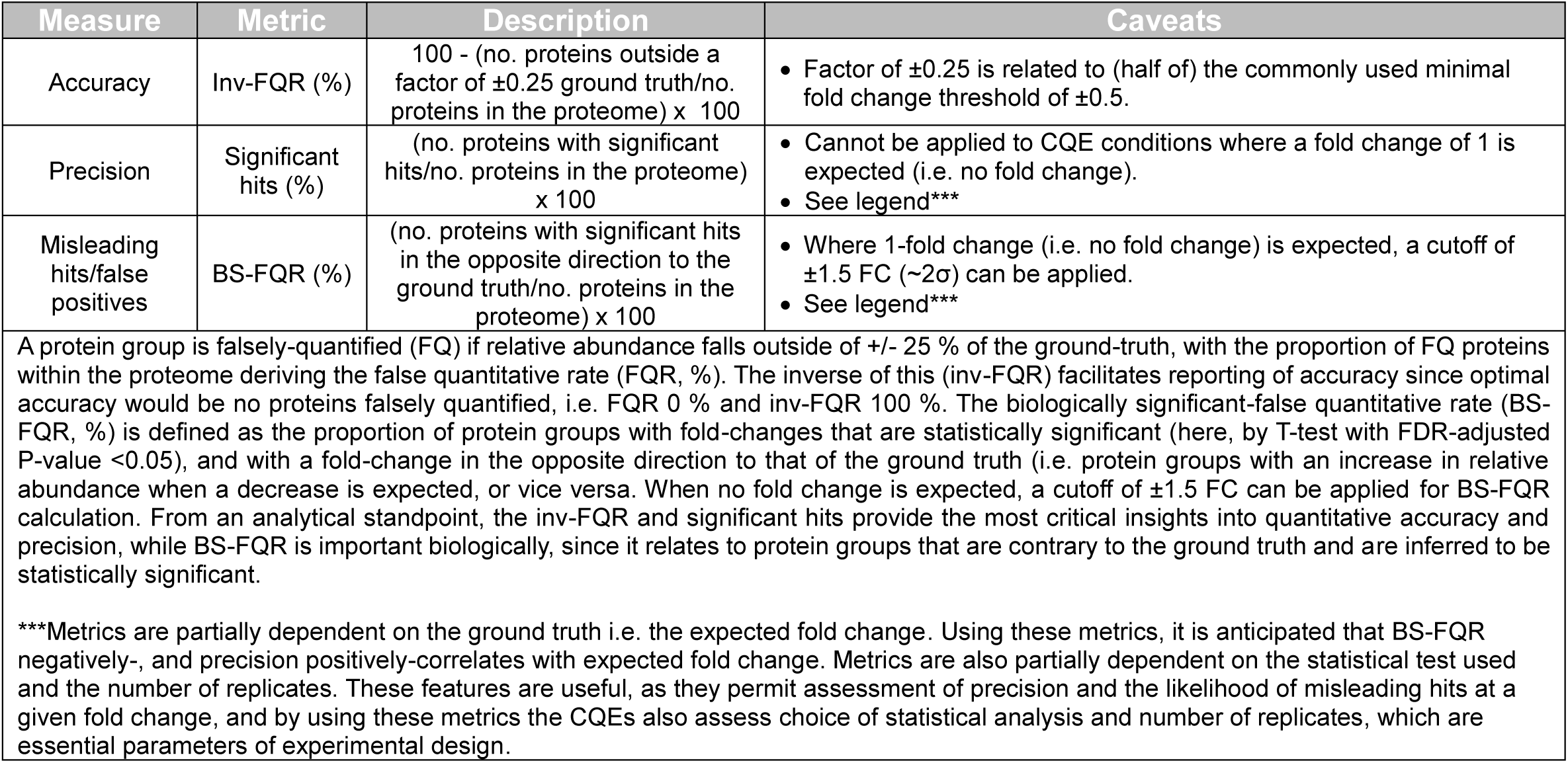
Description of CQE metrics.

Given the known composition of the mixtures, the optimal outcome would be characterised by human proteins with log₂ fold changes close to +1 and bacterial proteins close to −1, with no proteins exceeding the range of 1.5–2.5-fold change in the correct direction (corresponding to 100 % inv-FQR). In such an ideal case, all proteins would be statistically significant (100 % significant hits) with a BS-FQR of 0 %, and the number of quantified protein groups would be maximised.

### Recommendation: Perform Two-Species CQEs for DIA Parameter Optimization

When testing DIA acquisition parameters, a simple two-species CQE provides an effective means to assess quantitative performance while simultaneously capturing information on identification rates. In this study, human–bacterial protein mixtures were analysed using a 20 SPD (58-minute active gradient) method on the Evosep One coupled to a timsTOF HT operating in diaPASEF mode. MS1- and MS2-PASEF settings (16 windows, 2 frames) were held constant, while the trapped ion mobility spectrometry (TIMS) accumulation and ramp times (TART) were varied between 33, 66, and 100 ms, corresponding to duty cycles of approximately 0.6, 1.2, and 1.8 seconds, respectively. Protein digest loads of 50, 100, 500, and 1000 ng were tested (**Figure 1, Supplemental figure 1**). Post-processing involved filtering for proteins identified by at least two peptides per protein group, exclusion of putative contaminants, and statistical analysis using *limma* (13). The full output resulting from the analysis is provided (**supplemental table 4**).

At the highest load (1000 ng), approximately 10,000 unique proteins and 100,000–140,000 precursors were identified, with an average of 3.5–5.5 MS2 points per peak at full width half maximum (PPP-FWHM; **Supplemental figure 1**). Across all TART and load combinations, quantitation was robust (**Figure 1A**), with median fold changes of 0.97 ± 0.09 for human proteins and −0.80 ± 0.10 for bacterial proteins. Precursor and protein identifications increased with TART, whereas PPP-FWHM and MS2 signal decreased. Lower-abundance proteins were quantified with reduced precision and accuracy, as expected, and both metrics declined further with decreasing protein load (**Figure 1A**). Notably, very highly abundant proteins were less well quantified at higher TART values (**Figure 1B**), possibly due to space-charge effects within the TIMS cell or detector saturation.

Overall, the majority of quantitative metrics (**Figure 1**) indicated that a TART of 66 ms with a 1000 ng protein load offered the best performance, with minimal loss in total identifications compared to a 100 ms TART. Median normalisation distorted log₂ fold changes from the known ground truth (**Supplemental figure 2**) and was therefore deemed unsuitable for this dataset.

### Assessing Quantitative Efficacy Across the Entire Proteomics Pipeline Using Complex Proteomes

The two-species CQEs described above evaluate quantitative efficacy under conditions where most proteins change only modestly in abundance. As a result, conclusions drawn from these experiments may not be generalisable to scenarios involving more substantial proteome-wide changes. To address this, we designed an additional series of CQEs using three-species mixtures comprising human (HeLa), yeast (*S. cerevisiae*), and bacterial (*E. coli*) proteins (**Figure 2A**). The mixtures were prepared in defined ratios to produce large, modest, or small abundance changes affecting different proportions of the proteome. This design reflects a range of possible outcomes in biological experiments, enabling assessment of quantitative efficacy across varied scenarios and providing limited opportunities for false discovery rate (FDR) evaluation. Comparisons 1, 2, and 5 represent cases where only a small proportion of the proteome undergoes modest changes, which is most typical in biological studies. In contrast, comparisons 3, 4, and 6 represent extreme conditions where most or all proteins exhibit substantial abundance changes, as might occur when analysing distinct tissues, organelle-enriched fractions, or markedly divergent physiological states.

We also considered the potential influence of LC–MS hardware on quantitative performance. Accordingly, the three-species CQEs were performed on a timsTOF HT using the optimal acquisition settings identified earlier (66 ms TART, 1000 ng load) and compared to a method routinely utilised in our lab on an Ultimate 3000 RSLC-Nano coupled to an Orbitrap Exploris 480. In addition to sample composition, instrumentation, and acquisition parameters, post-processing decisions represent a critical factor in quantitative proteomics. Although numerous strategies exist for filtering and processing peptide-spectrum match (PSM) outputs, we initially focused on evaluating the effects of coefficient of variation (CV)-based filtering, exclusion of low-intensity precursors, minimum peptides-per-protein thresholds, and median normalisation on protein group–level quantitative efficacy. For each comparison (C1–C6), 32 distinct post-processing combinations were tested on both the timsTOF (**Supplemental figure 4**) and Orbitrap (**Supplemental figure 5**) datasets. Example outputs from three species CQEs following a post-processing pipeline (1-peptide per protein minimum, median normalised, contaminants excluded) datasets from the timsToF-HT and E480 are provided (**Supplemental tables 5-6**). The influence of post-processing parameters on quantitative performance was broadly consistent between scenarios with typical versus atypical proteome changes, allowing a single representative example to suffice for most recommendations.

### Recommendation: Median Normalisation is Usually Improving Data

Inspection of intensity and fold change profiles for the timsTOF HT (**Figure 2B, Supplemental figure 4**) and E480 (**Supplemental figure 5**) revealed that in typical comparisons (C1, C2, C5), fold-change distributions were already close to expected values without median normalisation, whereas in atypical comparisons (C3, C4, C6) observed fold changes were substantially skewed upward. Median normalisation exerted variable effects on quantitative efficacy across species and comparisons for both instrument platforms (**Supplemental figures 4 and 5**). In typical scenarios, modest decreases in quantitative performance were occasionally observed. For example, in C2 timsTOF data without additional processing (one peptide per protein), median normalisation reduced yeast proteome accuracy from 97% to 94%, precision from 91 % to 84 %, and increased BS-FQR from 0% to 0.1%, while having negligible impact on human proteome accuracy and BS-FQR (precision could not be meaningfully assessed for the human proteome in this comparison).

In contrast, for atypical comparisons, median normalisation consistently improved quantitative metrics across all species. In C3 timsTOF data, for example, median normalisation increased human proteome accuracy from near 0% to ∼50%, precision from ∼20% to ∼90%, and reduced BS-FQR from ∼7% to ∼0%. For yeast and bacterial proteomes in the same comparison, accuracy rose from ∼10% and ∼2% to ∼95% and ∼96%, respectively, while precision remained at ∼100% and BS-FQR near 0%.

Results from the Orbitrap dataset showed similar trends. In typical comparisons (C1, C2, C5), increasing the minimum peptide requirement generally improved quantitative metrics, with this effect being more pronounced when median normalisation was not applied.

Coefficient of variation (CV)-based filtering also enhanced quantitation in these cases, while median normalisation had little effect. For atypical comparisons (C3, C4, C6), however, median normalisation was critical. In C4, yeast protein precision dropped below 5% without normalisation, while in C3, median normalisation increased human protein precision from ∼85% to >98% and accuracy from <5% to substantially higher values. BS-FQR was also greatly reduced; in C3, human BS-FQR decreased from 1.5% to <0.1%, and in C4, absence of median normalisation resulted in yeast BS-FQR values exceeding 90%. Interestingly, in C4, increasing the minimum peptide requirement slightly reduced yeast protein accuracy and worsened BS-FQR despite median normalisation.

### Recommendation: Exercise Caution in Quantitative Efficacy for Atypical Comparisons Where the Majority of the Proteome Changes

Atypical comparisons (C3, C4, and C6) exhibited substantial deviations from the expected ground-truth fold changes. Prior to applying quantitation metrics or filtering parameters, careful consideration of raw data processing is essential, particularly regarding the search strategy and search space. In these experiments, protein and precursor information were generated using the open-source software DIA-NN (version 1.9), which applies cross-run normalization by default.

For the initial analysis, all raw files, including both typical and atypical comparisons, were processed together in a single search. To assess the impact of search scope and cross-run normalization on quantitative metrics, separate searches were also performed for each comparison individually. Quantitative metrics for these approaches are shown in **Figure 3**. Overall, searching atypical experiments separately generally decreased quantitation performance, resulting in lower accuracy and higher BS-FQR values. This suggests that for experiments in which a large portion of the proteome is changing, a broader search space may be beneficial.

For typical experiments, separate searches usually improved precision, but this came at the expense of accuracy, though BS-FQR values were improved for C1 and C2. These observations indicate that the effect of search space on quantitative metrics is largely experiment dependent. Interestingly, when atypical experiments were processed collectively, median normalization was critical for accurate quantitation (**Figure 3**). However, when these experiments were searched individually, median normalization had minimal effect. This highlights that in scenarios where median normalization is inappropriate, such as atypical experiments with widespread proteome changes, independent searches without median normalization provide a more suitable and accurate approach, as median normalization assumes that the majority of the proteome remains unchanged.

### Recommendation: Precursor Intensity Filtering Offers No Quantitative Benefit

In order to test if low-intensity precursors affect the quantitation negatively, precursors were filtered by excluding the lowest 1% by mean intensity (calculated across all replicates within the comparison) for each comparison. Across all comparisons and post-processing conditions for both timsTOF and Orbitrap datasets (**Supplemental Figures 4** **and 5**), this filtering approach was occasionally mildly detrimental to identification rates and had negligible effects on quantitative efficacy metrics. In the typical comparison C1 (median normalisation only, one peptide per protein, timsTOF data), the number of protein groups decreased slightly from 6695, 2315, and 1995 to 6695, 2311, and 1995 for human, yeast, and bacterial proteins, respectively. Corresponding accuracies declined marginally from 99.49 %, 83.55 %, and 98.17 % to 98.49 %, 83.49 %, and 98.12 %, while precision (for yeast and bacterial proteins) changed from 97.71 % and 99.78 % to 97.66 % and 99.78 %. BS-FQR values remained unchanged.

**Figure 4.**
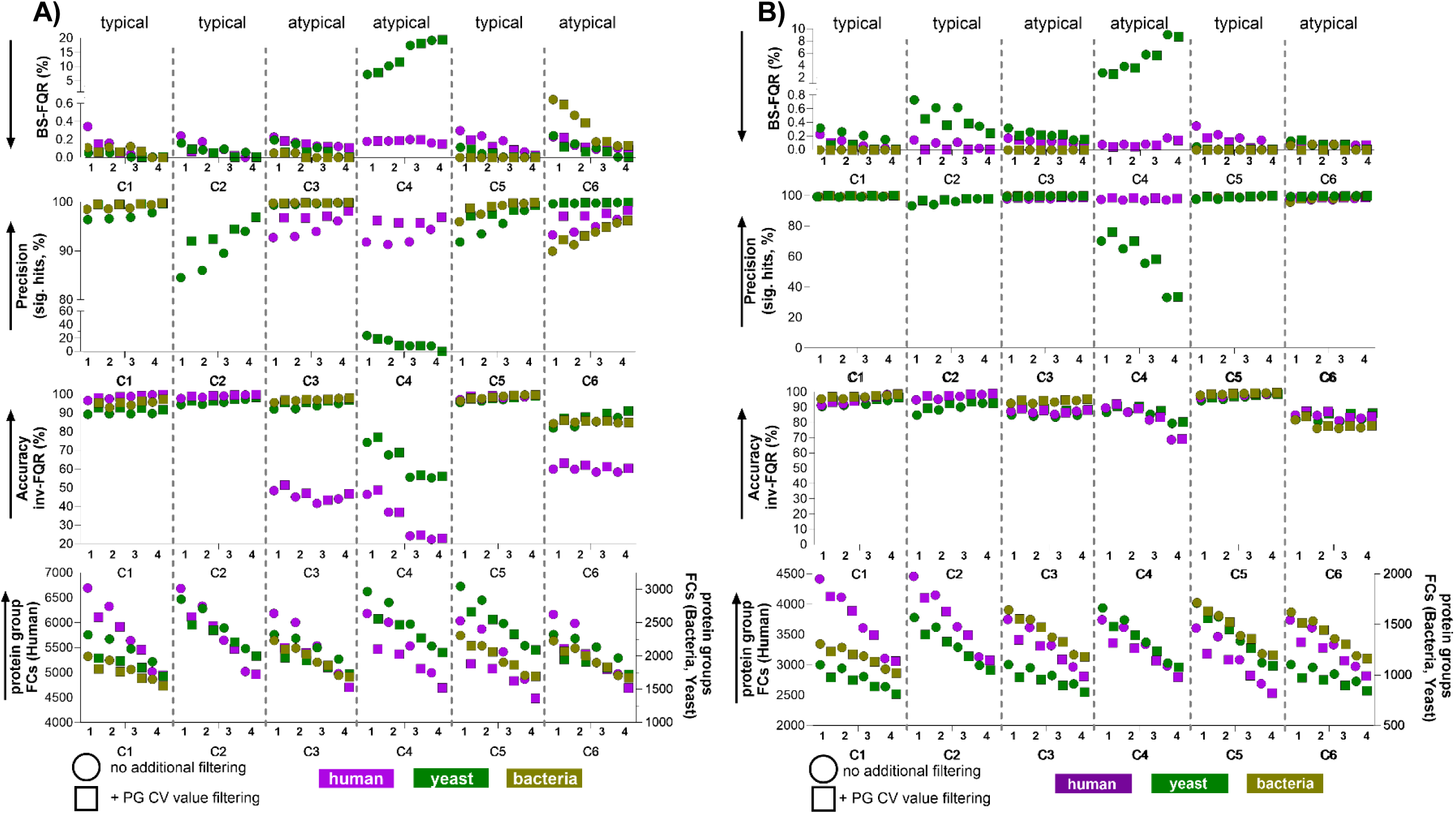
Filtering for peptide numbers per protein and protein CV have notable impacts on ID rates and quantitative efficacy in typical and atypical proteomics experiments. A) timsTOF-HT B) Orbitrap Exploris 480. Data shown underwent median normalisation, protein present in 3/5 replicates in both conditions, ± protein group CV filtering (square and circle symbols) and variable filtering by minimum-number of peptides (numbers on x-axis). Data derived from 5 replicates.

**Figure 5.**
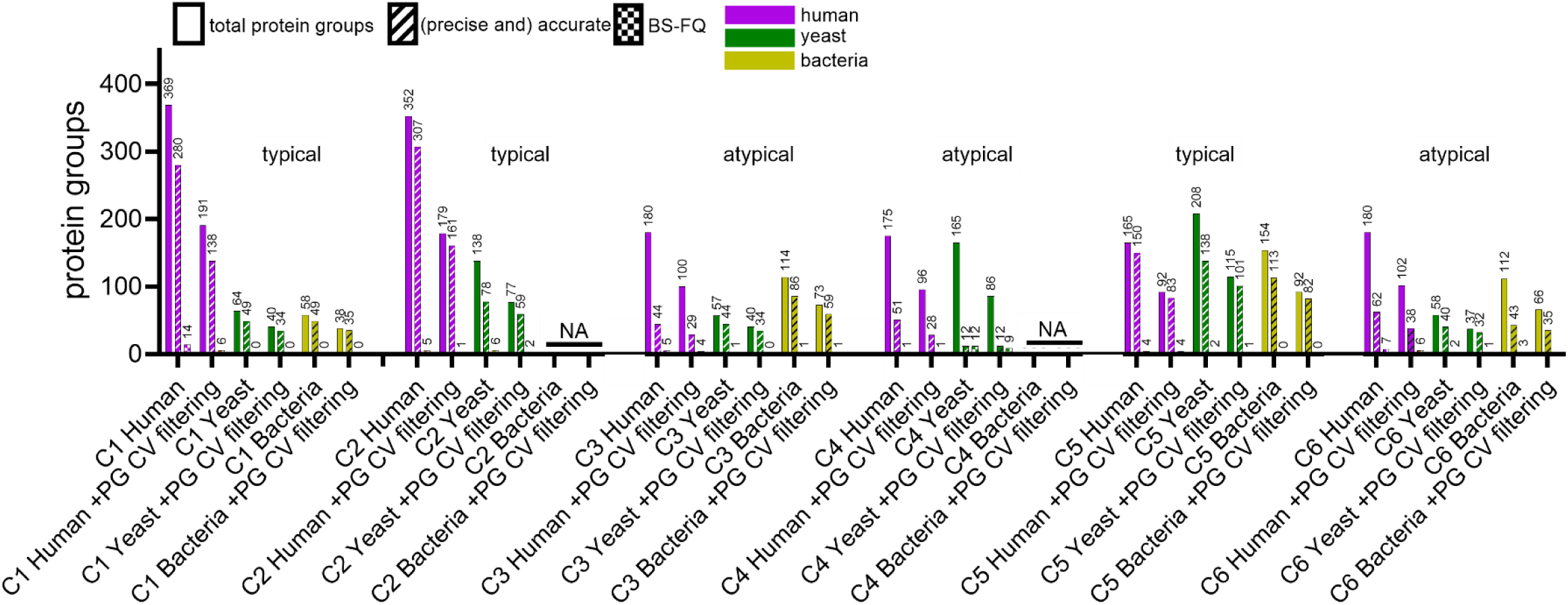
Single-peptide proteins and quantitative efficacy. Protein groups identified and quantified by a single peptide in the comparisons derived from the timsTOF-HT dataset (median normalised, ± protein group CV filtering) were plotted for each comparison, alongside the total number, those that are accurate (and precise where appropriate), and those that contribute to BS-FQR. Data are derived from five replicates.

In the atypical comparison C6, protein group counts decreased minimally from 6165, 2319, and 2230 to 6164, 2319, and 2228 for human, yeast, and bacterial proteins, respectively. Accuracy values shifted from 59.90%, 81.93%, and 84.26% to 59.23%, 82.23%, and 83.89%, while precision changed from 93.20%, 99.66%, and 90.09% to 93.07%, 99.66%, and 90.53%. BS-FQR values remained constant with the exception of bacterial proteins, which showed a slight decrease from 0.058% to 0.054%. Similar patterns were observed in the Orbitrap dataset, where precursor intensity filtering likewise had negligible effects on quantitative performance.

### Recommendation: While current proteomics pipelines display high quantitative efficacy, the most critical factors are low protein group coefficient of variation and high number of peptides-per protein

Next we tested if proteins with high coefficient of variation (CV) or few quantified peptides were a source of inaccuracy. Protein groups were filtered CV, requiring a CV <20 % in each condition of a given comparison for inclusion. Additionally, proteins were filtered to include only those quantified by a minimum number of peptides, with thresholds ranging from one to four peptides per protein tested. While both CV- and peptide-number-based filtering predictably reduced total protein group identifications, substantial improvements in quantitative efficacy were observed across all comparisons for both timsTOF and Orbitrap datasets (**Figure 4**), with the exception of C4, which is discussed separately. Remarkably, although the absolute number of protein group identifications differs between the Orbitrap and timsTOF systems, trends in quantitative efficacy metrics were highly consistent, particularly for typical comparisons.

In the typical comparison C1, applying a minimum threshold of four peptides per protein reduced human protein group counts by approximately 20 % relative to one-peptide filtering (timsTOF: 6107 to 5018; Orbitrap: 4417 to 3413), yet quantitative efficacy improved substantially. For human proteins, accuracy increased from 96.6% to 99.7% on the timsTOF and from 91.1% to 98.2% on the Orbitrap, while BS-FQR decreased from 0.34 % to 0% and from 0.22 % to 0.02%, respectively. Similarly, in the atypical comparison C6, human protein group counts decreased by 10–20% with four-peptide filtering (timsTOF: 5479 to 4977; Orbitrap: 3740 to 2974), but quantitative performance improved: accuracy increased from 96.6% to 99.7% (timsTOF) and 91.1% to 98.2% (Orbitrap), precision increased from 93.2% to 95.4% and 97.9% to 98.8%, and BS-FQR decreased from 0.23% to 0.1% and 0.09% to 0.07%, respectively.

As expected, proteins quantified by higher numbers of peptides generally exhibited lower CV, and therefore the effect of CV filtering on quantitative efficacy was less pronounced for proteins with ≥3–4 peptides than for those with one or two peptides. For example, in the typical comparison C1, BS-FQR for human proteins in the timsTOF dataset decreased substantially from 0.34% to 0.14% upon inclusion of CV filtering in the one-peptide dataset, and from 0.22% to 0.10% for the Orbitrap. In contrast, when the same comparison was filtered for four peptides per protein, BS-FQR was already 0% for the timsTOF without CV filtering and decreased from 0.02% to 0% for the Orbitrap.

### Recommendation: Single-Peptide Proteins Are Generally Well-Quantified and Rarely Cause Biologically Significant False Quantitation or Discovery

The data above showed that filtering out proteins with few quantified peptide, improved overall quantitation. However, next we wondered how good or bad proteins quantified with a single peptide were performing. We compared their precision, accuracy, and BS-FQR in each comparison (**Figure 5**), with and without protein group CV filtering in the timsTOF-HT dataset.

For typical comparisons (C1, C2, and C5), the majority of single-peptide protein fold changes are accurate, and very few represent biologically significant false quantitation. For example, in C5, of 165 single-peptide human proteins, 150 were accurate, and only 4 contributed to BS-FQR. For yeast and bacteria in the same comparison, 208 and 154 proteins were quantified, of which 138 and 113 were precise and accurate, and only 2 and 0 contributed to BS-FQR, respectively. While CV filtering modestly improved quantitative efficacy for single-peptide proteins, it did not fully eliminate false discoveries, and it also led to the loss of ∼50 % of accurate human proteins in typical comparisons (C1, C2, C5) and ∼20 % of accurate and precise proteins in atypical comparisons across human, yeast, and bacterial datasets.

Regarding false discoveries, a small number of *E. coli* proteins (≤10) were detected in the HYB 70:30:0 condition, where they should not have been present, leading to some *E. coli* protein groups in comparisons C2 and C4 (**Supplemental figure 6, Supplemental table 7**). Applying an additional filter to include only proteotypic precursors - precursors unique to a single protein - had minimal impact on these falsely discovered *E. coli* proteins. Notably, all of these proteins were quantified by 4–39 peptides across the comparison, indicating that removing single-peptide hits does not prevent these apparent false discoveries. Protein group CV filtering, however, removed most falsely quantified proteins, which were generally close to no-fold change (within ±1.5-fold) or changed in the expected direction for the comparison (**Supplemental figure 6 B**). In comparison C4, one *E. coli* protein remained significantly decreased beyond the ±0.5-FC cutoff (-2.26-fold change), regardless of CV filtering.

### Recommendation: Apply a Minimum Fold-Change Threshold of ±1.5 to Limit False Discoveries in DIA-LFQ

To prevent proteins with low quantitative accuracy being annotated as “significant”, a fold-change cutoff is often applied alongside appropriate statistical tests. Here, a ±1.5-FC threshold (0.75–1.5 fold) was used, although more stringent cutoffs could be applied if desired. In typical comparisons where the human proteome is expected to be unchanged, small fold changes were still observed for all quantified proteins, generally close to 1 (i.e., no true fold change; **Figure 2B**).

These minor, artefactual abundance shifts were measured consistently enough that a substantial fraction of proteins appeared statistically significant, despite stringent Benjamini-Hochberg adjustment of p-values. In C1 and C2 comparisons, for example, where human proteins stayed constant, over 50% of human proteins were classed significant by p-value alone (**Supplemental figure 7**). By applying the ±1.5-FC cutoff, inference of “biological significance” for these proteins is appropriately restricted, resulting in very low BS-FQR for human proteins (**Figure 4, Supplemental figure 4)**. Implementing this fold-change threshold also effectively mitigates the false quantification and discovery of proteins described in the previous section (**Supplemental figure 6**).

### Recommendation: Exclusion of Precursors with Low Points per Peak May Not Benefit Protein Group Quantitation, Though Effects Are Instrument- and Method-Dependent

A contentious parameter in untargeted proteomics is the minimum number of points per peak (PPP) required for acceptable quantitative efficacy, and whether protein group quantitation should rely on precursors with low PPP. To assess this, different PPP filters were applied to timsTOF and Orbitrap datasets. For each precursor, DIA-NN reports the full width at half maximum (FWHM) in minutes, allowing calculation of approximate PPP-FWHM based on the MS/MS cycle time (1.2 sec for timsTOF and 1.6 sec for Orbitrap). A range of PPP filtering parameters was tested (**Figure 6**).

**Figure 6.**
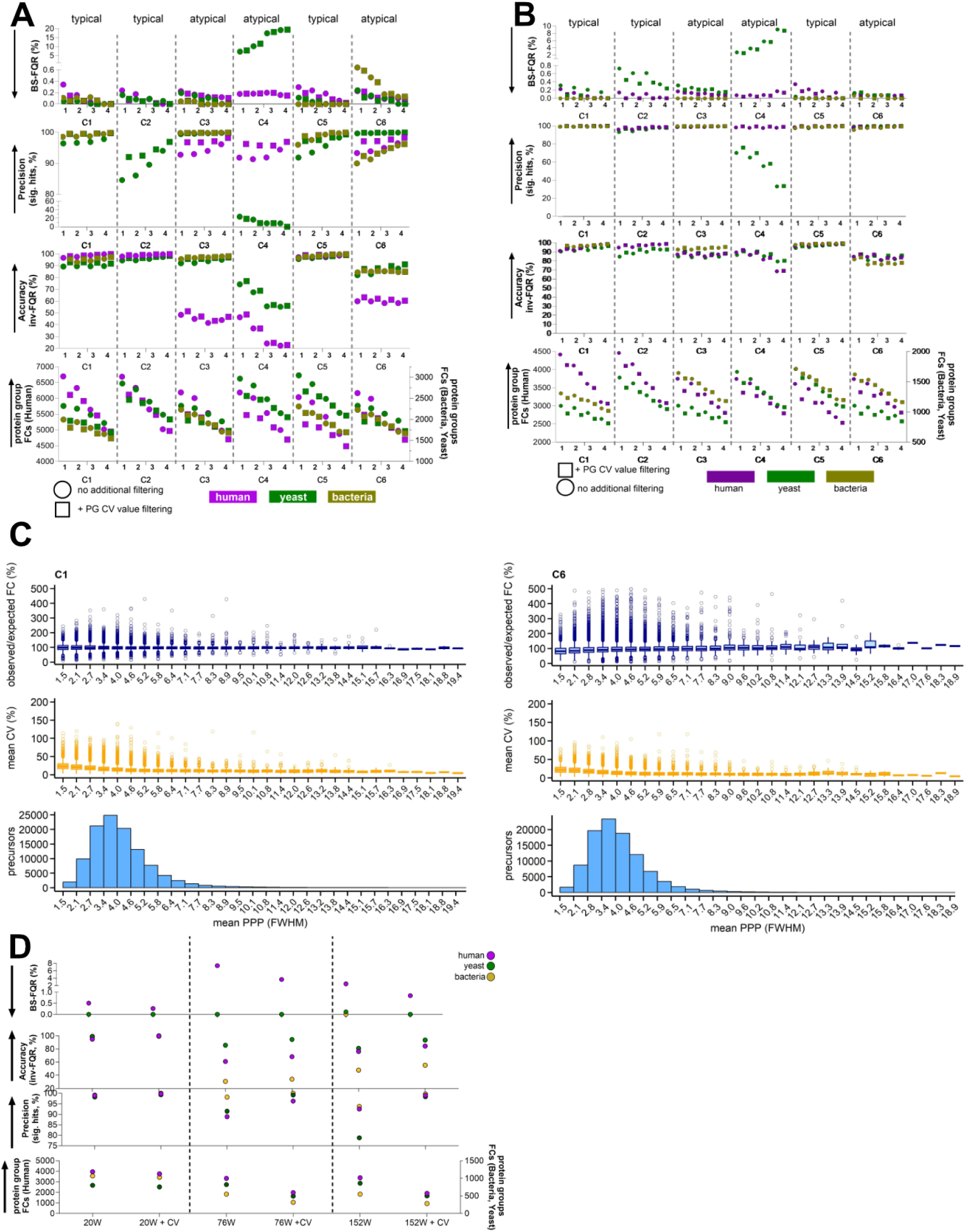
The impact of Precursor PPP filtering on Quantitative Efficacy. A) and B) timsTOF and Orbitrap-quantified protein groups subjected to different modes of PPP-filtering. (all) = global precursor filtering, (av.) = precursor filtering based on mean PPP-FWHM across all replicates from all conditions in a given comparison. C) timsTOF precursors (with no filtering) were placed into 30 bins based on their PPP, for each comparison. For each precursor bin, the number of precursors was plotted-lowest histogram, as was the median CV (%) and deviation from ground truth (observed/expected fold change, %)-middle and upper boxplot. Boxplots show median, quartiles, outliers. Data are derived from 5 replicates. D) Quantified protein groups using MS data acquired with different DIA window schemes (20, 76, and 152 windows), both with and without CV filtering.

For timsTOF data (**Figure 6A**), a global filter of 3 PPP-FWHM applied to all precursors before assembling protein groups and performing single-peptide-per-protein analysis unexpectedly had a mildly detrimental effect on most quantitative metrics across all species. In C1, human, yeast, and bacterial protein group fold changes decreased from 6695/2315/1995 to 6143/1962/1825, yeast/bacterial precision dropped from 96.4%/98.6% to 94.4%/97.7%, accuracy shifted from 96.6%/89.3%/92.6% to 95.1%/88.6%/92.9%, and BS-FQR slightly increased. The bacterial condition showed a minimal improvement (≤0.5%), but most metrics worsened. Missing value inspection revealed that filtering disproportionately removed precursors even within replicates of the same condition.

Filtering by *mean* PPP-FWHM across replicates (PPP(av.)), separately for each comparison, produced similar metrics to no filtering, though with slightly reduced protein group IDs (C1 FCs dropped from 6695/2315/1995 to 6411/2114/1915). Moderate PPP thresholds (3 PPP(av.)) had minimal effect, but stricter cutoffs (6 or 10 PPP(av.)) caused severe ID losses and worsened quantitative efficacy - especially in atypical comparisons -while increasing BS-FQR (e.g., C6 bacterial BS-FQR rose from 0.58% to 2.24% at 6 PPP(av.)). In the most stringent case, IDs dropped below 400 (human) and 200 (yeast/bacteria) for all comparisons.

Precursor PPP distribution analysis (**Figure 6C, Supplemental figure 8**) showed a modal average of 3.4–4.0 PPP-FWHM, with median precursor CVs ∼25% in most bins (rising to ∼50% at 1.5–2.1 PPP-FWHM). Despite this, observed/expected protein group fold changes were ∼100% across bins, suggesting that protein-level quantitation is largely independent of individual precursor PPP in the timsTOF datasets.

By contrast, Orbitrap E480 data benefited more from PPP filtering (**Figure 6B)**. A 3 PPP-FWHM threshold reduced IDs only slightly (3–4%), indicating most precursors already exceeded ∼8 PPP. Increasing to 6 PPP-FWHM caused larger ID losses (19–23%) but generally improved precision, accuracy, and BS-FQR (e.g., C4 yeast precision rose from 74% to 90%). Some exceptions occurred (e.g., C6 yeast precision dropped from 98% to 90%), indicating instrument- and experiment-specific effects.

The relatively limited impact of PPP filtering in timsTOF data can likely be attributed to previous careful DIA method optimisation, balancing PPP, cycle time, and reproducibility to yield consistently low CVs. To test the deliberate changes in PPP, we analysed C1 samples with three DIA schemes on the Exploris 480:

- **152 DIA windows** with 2 m/z overlap (cycle time 12.6 s, ∼3 PPP-FWHM)
- **76 DIA windows** with no overlap (cycle time 6.3 s, ∼4.5 PPP-FWHM)
- **20 DIA windows** with 12 m/z overlap (cycle time 1.6 s, ∼5.9 PPP-FWHM)

Quantitative efficacy metrics (**Figure 6D**) show that the 20-window scheme delivered the strongest performance across all measures. Methods with fewer PPP (e.g., 152-window) suffered greater ID losses upon CV filtering, likely due to higher precursor/protein detection variability. Comparing the overlapping 152-window to the non-overlapping 76-window scheme suggested that window overlap can modestly improve accuracy and BS-FQR when combined with CV filtering.

Overall, PPP filtering effects are both instrument- and method-dependent: there was minimal benefit (and some detriment) for the timsTOF when acquisition was already optimised, but potential improvements for Orbitrap datasets at moderate PPP thresholds.

### Recommendation: Not only identification rates, but also quantitative efficacy must be considered for low-input or higher-throughput proteomics approaches

While most current proteomics experiments utilize hundreds of nanograms to micrograms of material at modest or low throughput, the field is trending toward shorter run times per sample and smaller amounts of input material. In parallel, new hardware (4, 14) and novel or adapted acquisition modes are being reported. Most studies emphasize protein identification rates, but quantitative efficacy must also be evaluated before widespread adoption of these methods, and any limitations must be clearly defined.

We applied this principle when testing high-sensitivity methods on the timsTOF-HT. Initially, 250 pg of tryptic HeLa digest was analysed by diaPASEF using 4, 6, or 16 frames, with TARTs of 100, 200, 300, or 400 ms, and either standard (m/z 300–1200) or focused (m/z 450–950) mass ranges, at a throughput of 40 SPD (**Supplemental figure 9**). Overall, 200– 300 ms TART, 4W methods were most optimal based on observed metrics (protein group IDs, protein group CV, and mean precursor PPP), achieving ∼3000 protein groups, protein group CV <15%, and ∼2.5–3 mean PPP-FWHM. While these metrics are good given that the hardware is not optimized for single-cell experiments, quantitative efficacy cannot be inferred from these metrics alone.

Therefore, a single CQE was performed using 250 pg of the defined species tryptic digest (C1, H:Y:B 70/20/10 → 70/10/20), testing key post-processing parameters (**Figure 7**): Minimum 1- or 4-peptides-per-protein filtering, with or without protein group CV filtering, and re-evaluating precursor intensity and PPP filtering under these high-sensitivity conditions.

**Figure 7.**
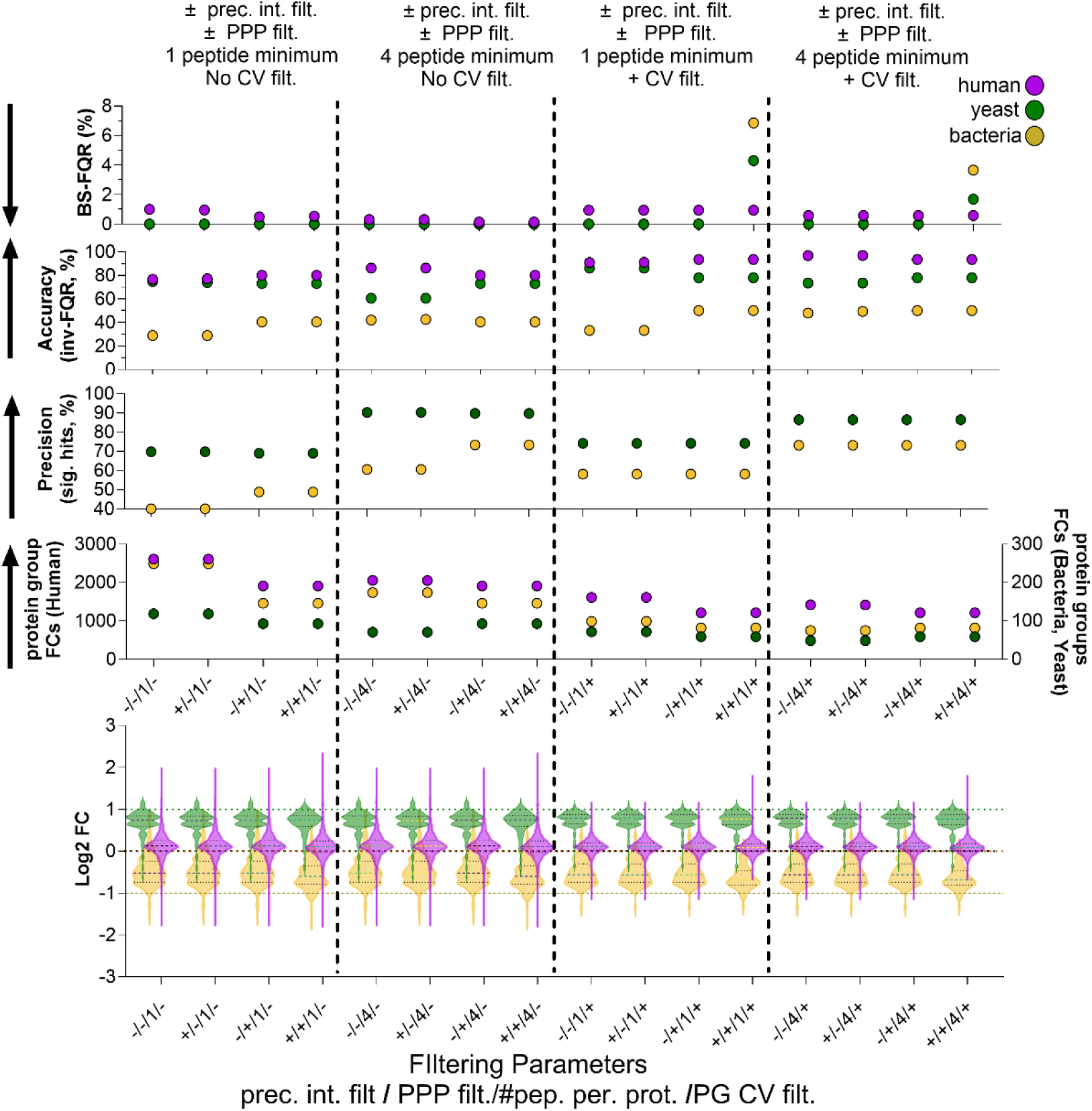
Quantiative efficacy of the timsTOF-HT at single-cell level sensitivity. Data underwent median normalisation, protein present in ≥3/5 replicates in both conditions. Then as indicated, ±precursor intensity filtering (bottom 1 % excluded), ± PPP-filtering (mean PPP-FWHM ≥3 across all replicates from all conditions in a given comparison), minimum-number of peptides, and ± protein group CV filtering (square and circle symbols). Data derived from 5 replicates.

Across all processing conditions, log2 fold-change distributions suggested compression, particularly for bacterial proteins, with observed fold changes skewed toward zero (Y; B log2FC medians ∼0.75; –0.55). This was reflected in quantitative metrics: with minimal filtering, bacterial precision was ∼40% and accuracy ∼30%, while yeast and human performed better (∼70% precision, ∼75% accuracy). BS-FQR remained low across species (H;Y;B = 1%;0%;0%).

Filtering by ≥4-peptides-per-protein generally improved quantitative efficacy. Compared with ≥1-peptide-per-protein filtering, yeast and bacterial precision increased from 69.8%;40.1% to 90%;61%, H;Y;B accuracy from 76.6%;74.8%;28.9% to 86.0%;60.6%;42%, and BS-FQR decreased slightly from 1%;0%;0% to 0.3%;0%;0%. Protein group CV filtering also improved quantitative metrics, while precursor intensity filtering had minimal impact. Points-per-peak (PPP) filtering provided moderate benefit, improving precision (Y;B 69.8%;40.1% → 69.0%;48.9%) and accuracy (H;Y;B 76.6%;74.8%;28.9% → 80.1%;73.1%;40.4%), with minor effects on BS-FQR. Interestingly, applying all high-stringency filters together (PPP, intensity, and protein group CV) increased BS-FQR for yeast and bacterial proteomes, indicating overly stringent filtering can be counterproductive.

Overall, the CQE revealed that this diaPASEF scheme on a timsTOF-HT for single-cell and ultra-high-sensitivity proteomics exhibits fold-change compression, reducing quantitative accuracy for proteins with differential abundance. Nevertheless, the low BS-FQR (<1%) indicates that proteins identified as significantly changing (±1.5-FC) are likely true positives, especially when quantified by ≥4 peptides and/or precursors ≥3 average PPP. Importantly, ≥4-peptides-per-protein filtering alone enhanced quantitative efficacy more effectively than ≥1-peptide-per-protein filtering combined with PPP filtering. When PPP filtering is applied, high peptide-per-protein filtering should also be used to maximize quantitative efficacy without further loss of protein fold-change. For example, moving from ≥1-peptide-per-protein + PPP filtering to ≥4-peptides-per-protein + PPP filtering increased Y;B precision from 69%;40.2% to 89.8%;73.3%, maintained H;Y;B accuracy at 80.1%;73.1%;40.4%, and improved BS-FQR from 0.5%;0%;0% to 0.14%;0%;0%, while protein group fold-changes remained constant (1915/93/146).

## Discussion

The widespread adoption of data-independent acquisition (DIA) on modern mass spectrometers has dramatically improved throughput and sensitivity, enabling experiments such as single-cell proteomics that seemed unattainable only a few years ago. While much of the field has focused on maximizing the number of identified proteins, less attention has been paid to the quantitative accuracy of these approaches.

Controlled quantitative experiments (CQEs) have proven invaluable for benchmarking emerging instruments and methods (7–9, 11, 15). In this study, we employed a range of CQEs to systematically evaluate acquisition parameters and data-processing strategies, and we present a set of practical recommendations for the community.

First, we advocate the use of simple two- or three-species CQEs to optimize any novel DIA (or DDA) workflow. This design provides a robust framework for balancing protein identifications with quantitative accuracy, and it is particularly informative across different sample loads, where fill times may need adjustment. As expected, lower sample inputs reduced protein identifications and compromised quantitation. Nevertheless, even at 250 pg loading, overall proteome-level quantitation remained robust, although with compressed ratios and decreased precision compared to standard protein inputs. These observations are directly relevant to the rapidly expanding field of single-cell proteomics, where such trade-offs are unavoidable and must be carefully considered.

For most conventional proteomics applications -where only a small subset of proteins changes abundance -standard workflows already perform well. However, atypical experiments involving large-scale proteome changes may require tailored approaches. Our data suggest that normalization, while generally beneficial, can in some cases introduce distortions, particularly when combining datasets from typical and atypical experiments.

CQEs also enable researchers to assess which instrument platform is most suitable for a given application. By comparing two widely used instruments, we identified performance differences that highlight complementary strengths and weaknesses. For instance, the rapid scan speed of TOF analysers appears to mitigate quantitation issues associated with reduced points per peak, in contrast to slower Orbitrap analysers. Conversely, we observed ratio compression for high-abundance precursors on the timsTOF, potentially arising from space-charge effects or detector saturation. Importantly, this effect had minimal impact at the protein level, as high-abundance proteins were typically quantified by multiple peptides.

In addition, CQEs are a powerful tool for evaluating false discovery rates (FDR) of search algorithms. Across our analyses, we detected very few false-positive protein identifications.

In one stringent comparison (C4), where no *E. coli* should have been detected in one condition, only a small number of *E. coli* proteins were identified, well below the nominal 1% FDR threshold. This suggests that current tools may be even more stringent than their stated FDR estimates.

Overall, our results reinforce that established proteomics workflows already deliver excellent quantitation, low false quantitation rates (FQR), and robust FDR control. Even proteins supported by a single peptide were generally well quantified, underscoring the reliability of modern DIA-based proteomics. These findings should reassure the community that proteomics is a powerful and trustworthy quantitative approach, capable of driving biological and medical discoveries. This is particularly timely as non-mass spectrometry-based platforms are being marketed as high-throughput proteomics solutions, often without equivalent benchmarking against established methods.

Finally, we summarize our recommendations in **Table 2**, which outline best practices for acquisition, processing, and evaluation of DIA data. We hope these guidelines will assist researchers in designing, executing, and interpreting proteomics experiments, and will promote robust and reproducible applications of DIA across the field.

**Table 2.** Recommendations for Quantitative Proteomics using DIA-LFQ.

| Table 2. Recommendations for Quantitative Proteomics using DIA-LFQ |
| --- |
| 1) While current proteomics pipelines display high quantitative efficacy, the most critical factors are low protein group coefficient of variation and high number of peptides-per protein. |
| 2) Apply a Minimum Fold-Change Threshold of $\pm 1.5$ to Limit False Discoveries in DIA-LFQ |
| 3) Not only identification rates, but also quantitative efficacy must be considered for low-input or higher-throughput proteomics approaches. |
| 4) Single-peptide proteins are generally well-quantified and rarely cause biologically significant false quantitation or discovery. |
| 5) Exclusion of precursors with low points per peak may not benefit protein group quantitation, though effects are instrument- and method-dependent. |
| 6) Precursor intensity filtering offers no quantitative benefit. |
| 7) Median normalisation is usually improving data. |
| 8) Perform Two-Species CQEs for initial DIA parameter optimization. |
| 9) Exercise caution when considering quantitative efficacy for atypical comparisons, where the majority of the proteome changes. |

## Acknowledgements

This research was partly funded by a Wellcome Trust Investigator Award to MT (215542/Z/19/Z) and a Wellcome Trust multi-user equipment grant (212947/Z/18/Z). We would like to thank all members of the Laboratory for Biomedical Mass Spectrometry for helpful discussions. We would like to thank Dr Katharina Trunk for providing the yeast lysates.

## Abbreviations

AGC: (Automatic gain control)
BS-FQR: (Biologically significant false quantitation rate)
CQE: (Controlled quantitative experiment)
CV: (Coefficient of variation)
DDA: (Data dependent acquisition)
DIA: (Data independent acquisition)
FDR: (False discovery rate)
FQR: (False quantitation rate)
FWHM: (Full width half maximum)
IAA: (Iodoacetamide)
ID: (Identification)
LC: (Liquid chromatography)
LC-MS/MS: (Liquid chromatography mass spectrometry)
LFQ: (Label free quantification)
MS: (Mass spectrometry)
(PSM): Peptide spectrum match
QC: (Quality control)
SDS: (Sodium doecyl-sulfate)
TART: (tims accumulation and ramp time)
TCEP: (Tris(2-carboxyethyl)phosphine hydrochloride)
TDC: (Target decoy competition)
TEAB: (Triethylammonium bicarbonate buffer)
TIMS: (Trapped ion mobility spectrometry)
TOF: (Time of flight)

