## Supplemental Figures for "Recommendations for Quantitative Data-Independent Acquisition (DIA) Proteomics using Controlled Quantitative Experiments (CQEs)"

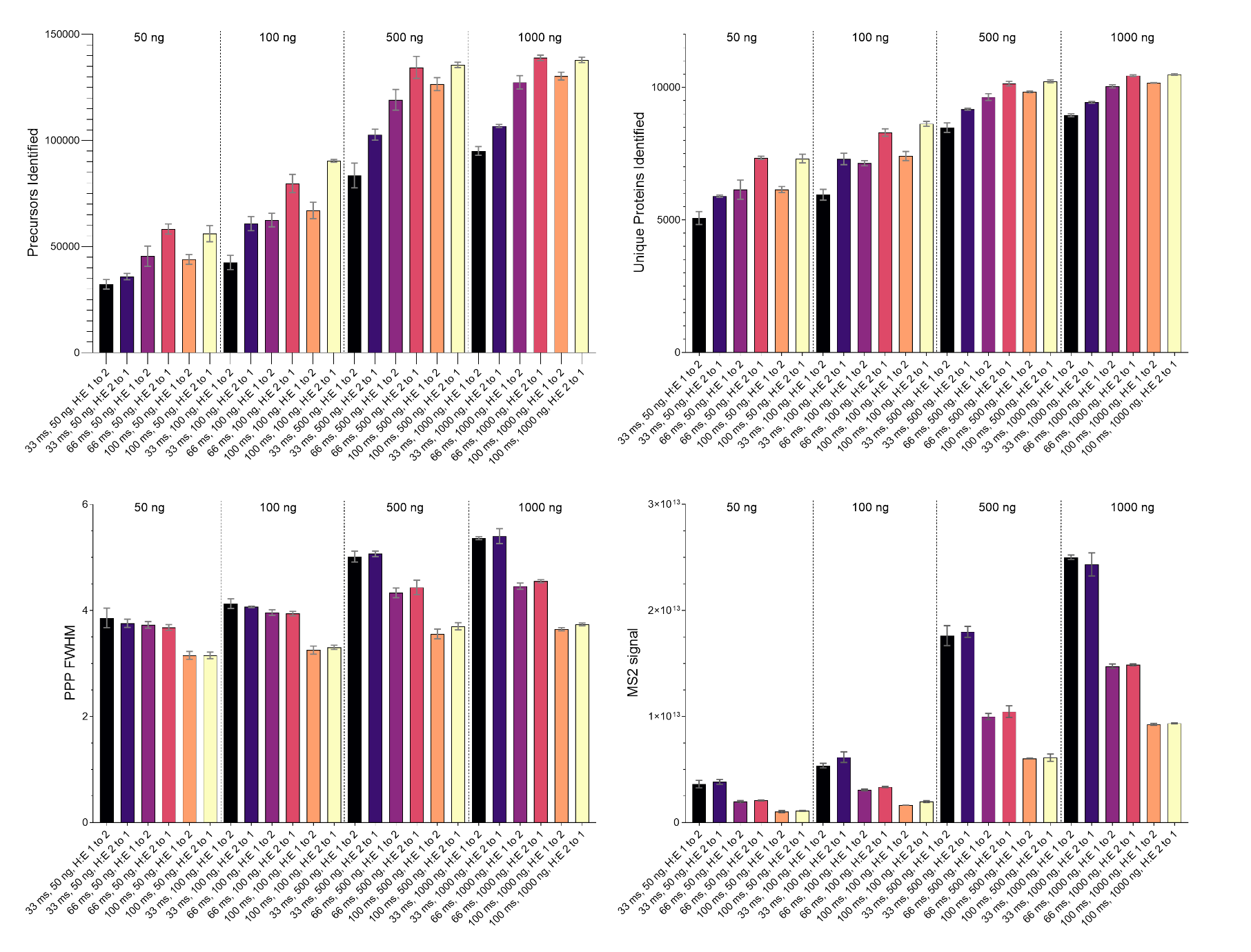
**Supplemental figure 1. Summary quality control metrics for all load and TART conditions.** Data shown are the mean of three replicates, error bars = standard deviation. PPP FWHM = points per peak, full width at half maximum.

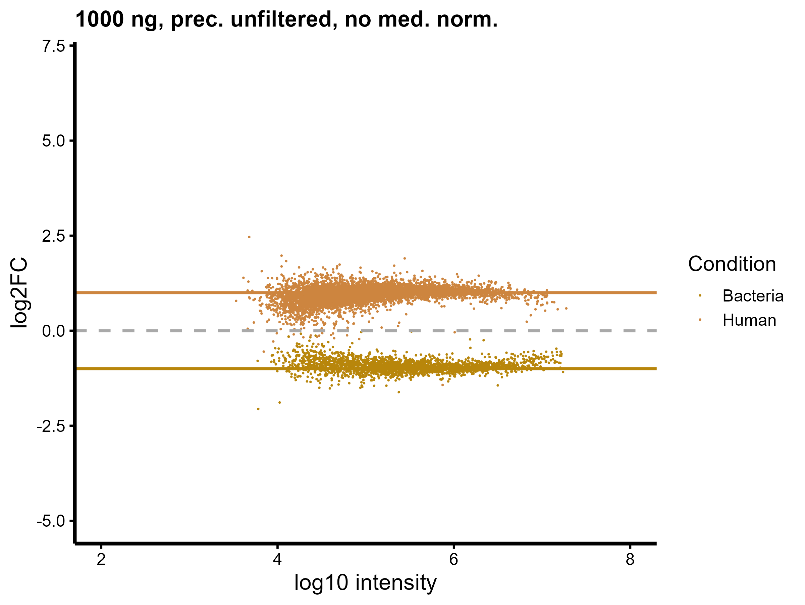

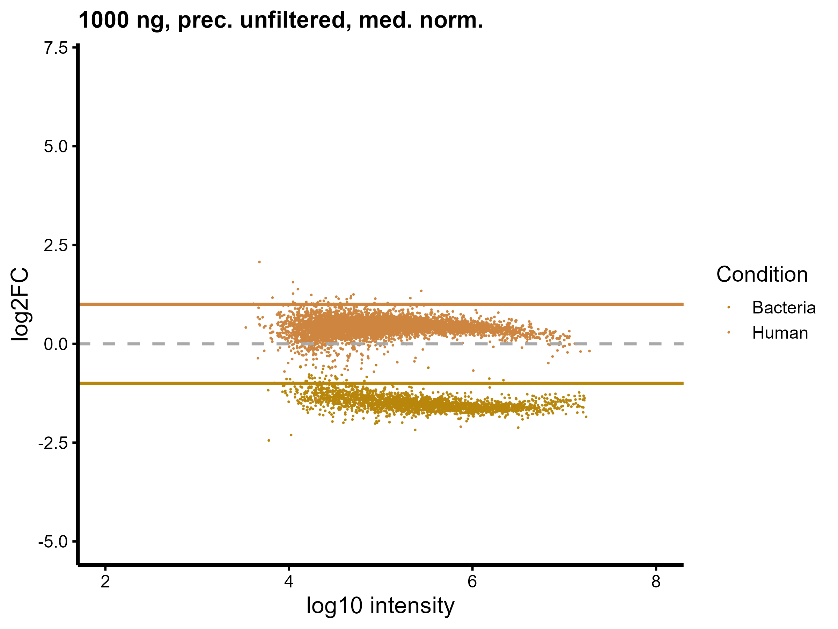
**Supplemental figure 2. Normalisation skews data away from the ground truth when the majority of the proteome is expected to change in abundance**. Distribution of (log10) protein group intensities against (log2) fold change for all conditions. Terracotta = Human, Khaki = Bacteria. Data shown are derived from three replicates.

Not-normalised

Median-normalised

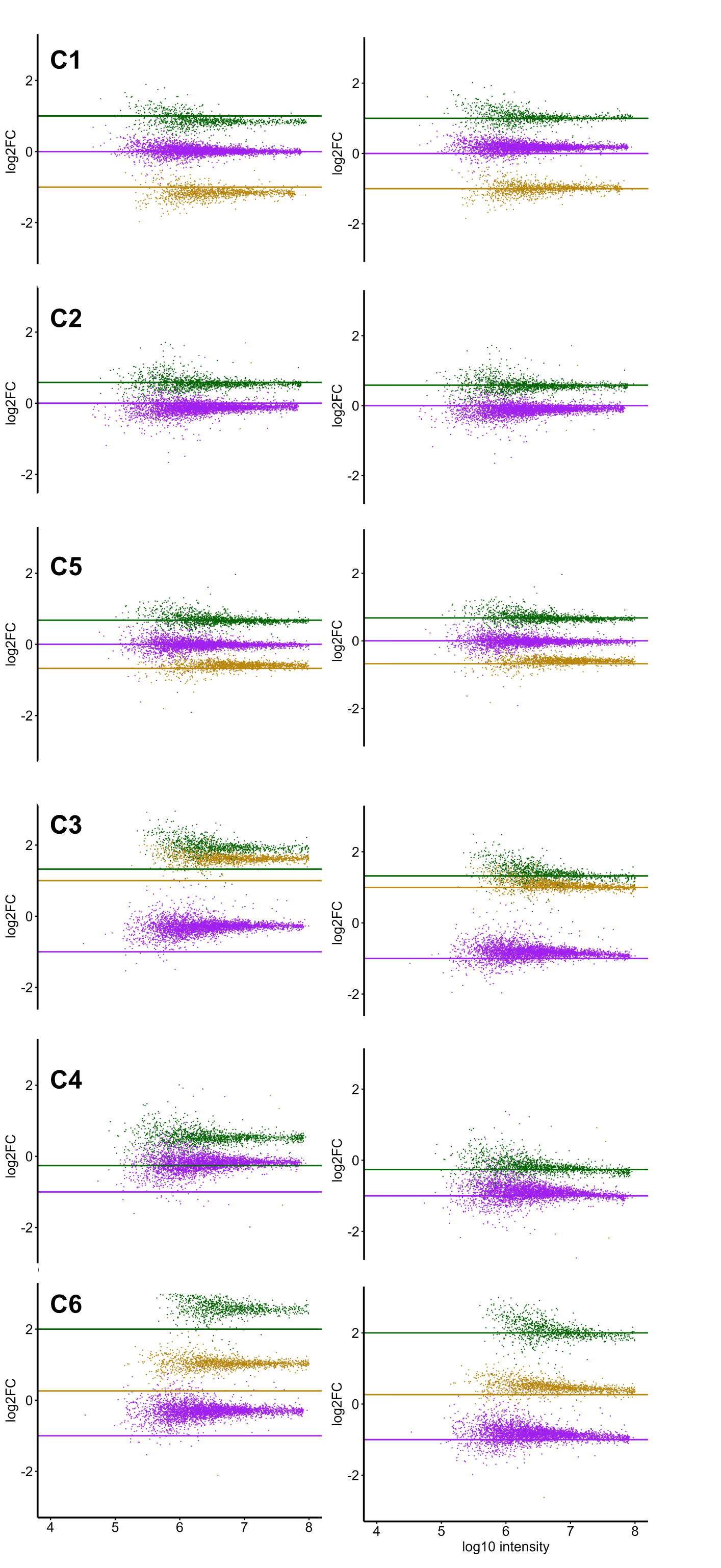

**Supplemental figure 3.** Median normalisation alters distribution of measured protein group FC for E480 data. FC shown here is the default post-processing pipeline (2 peptides per protein, protein present in each condition)

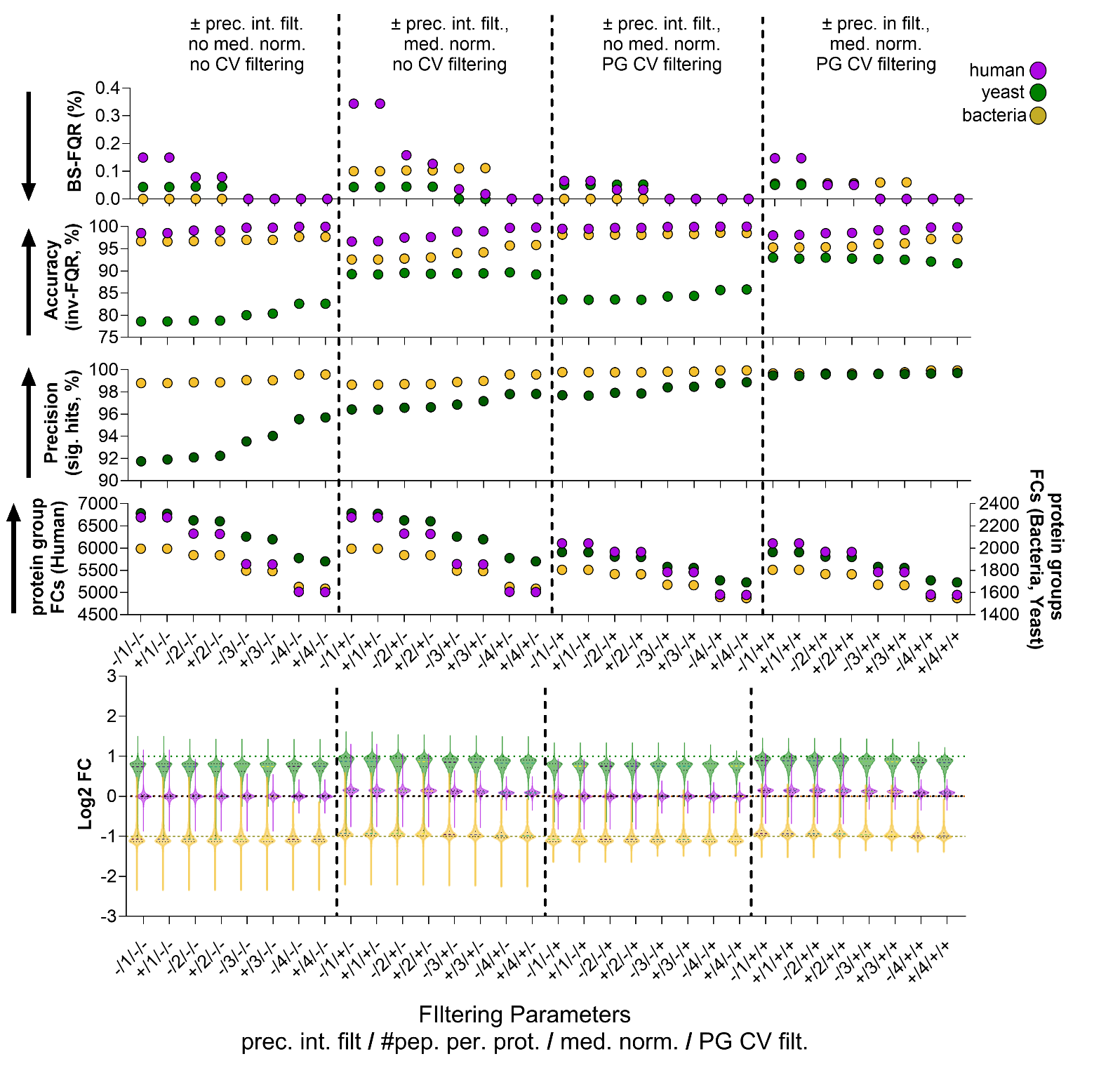

**C1**

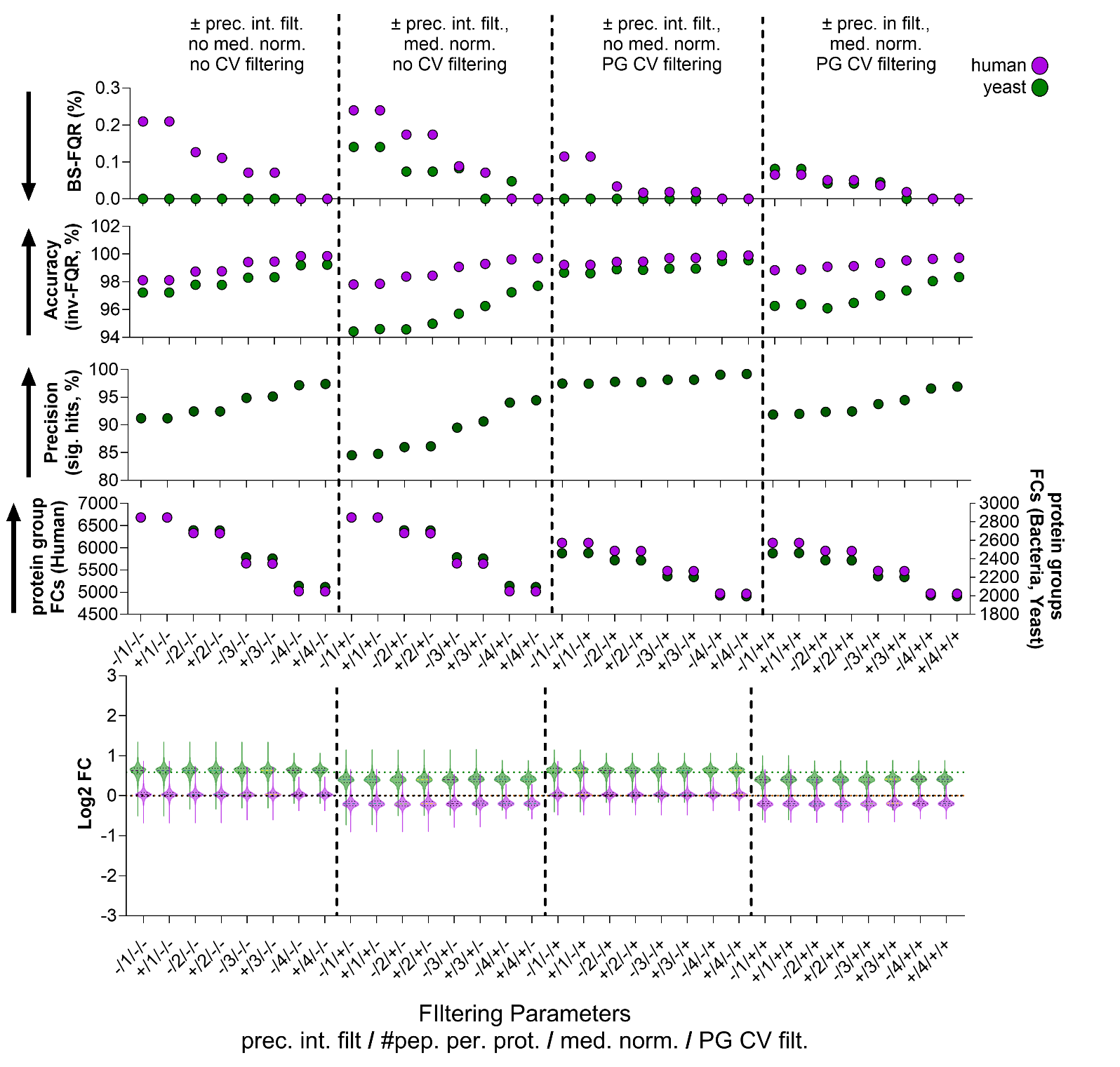

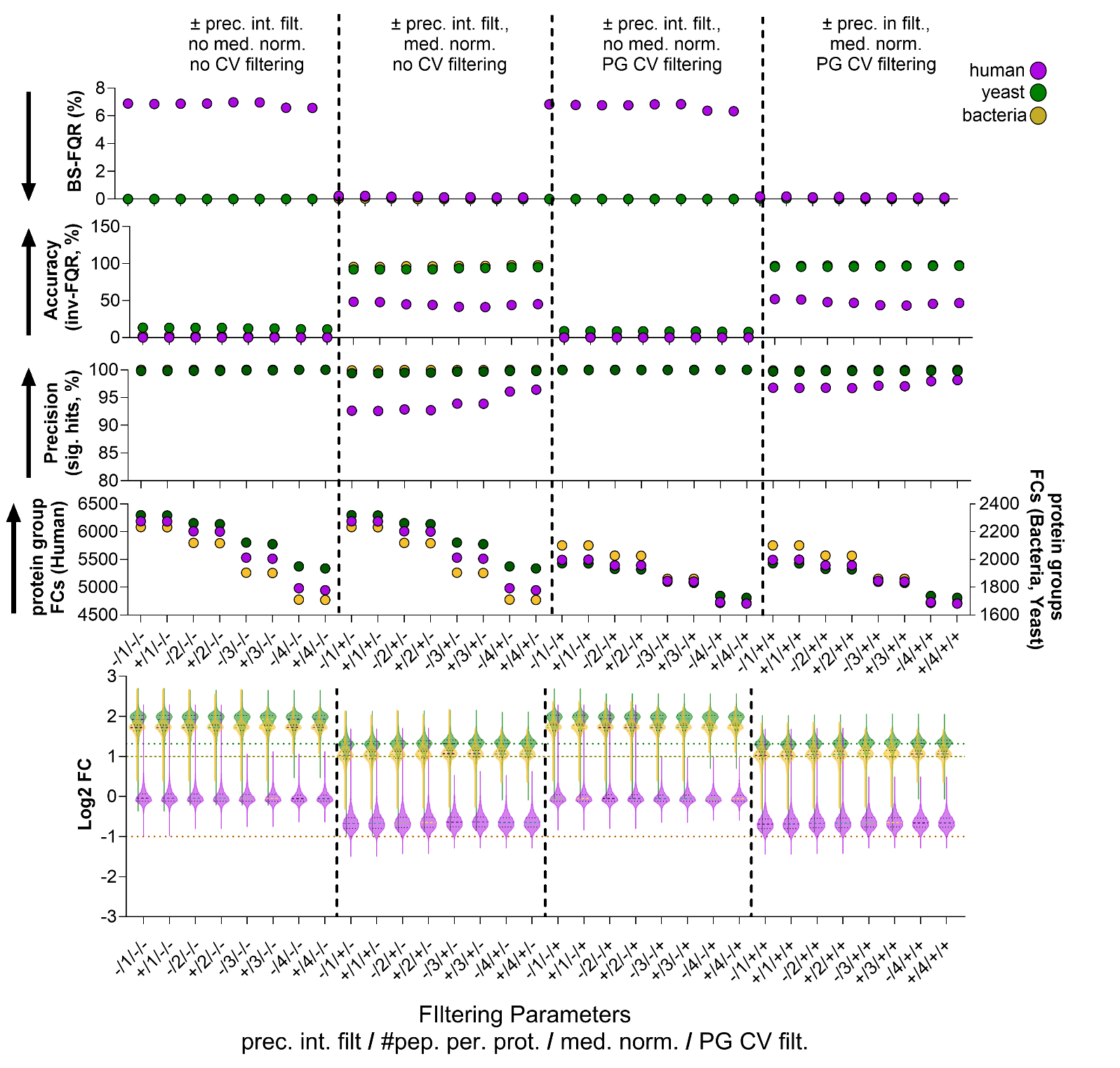

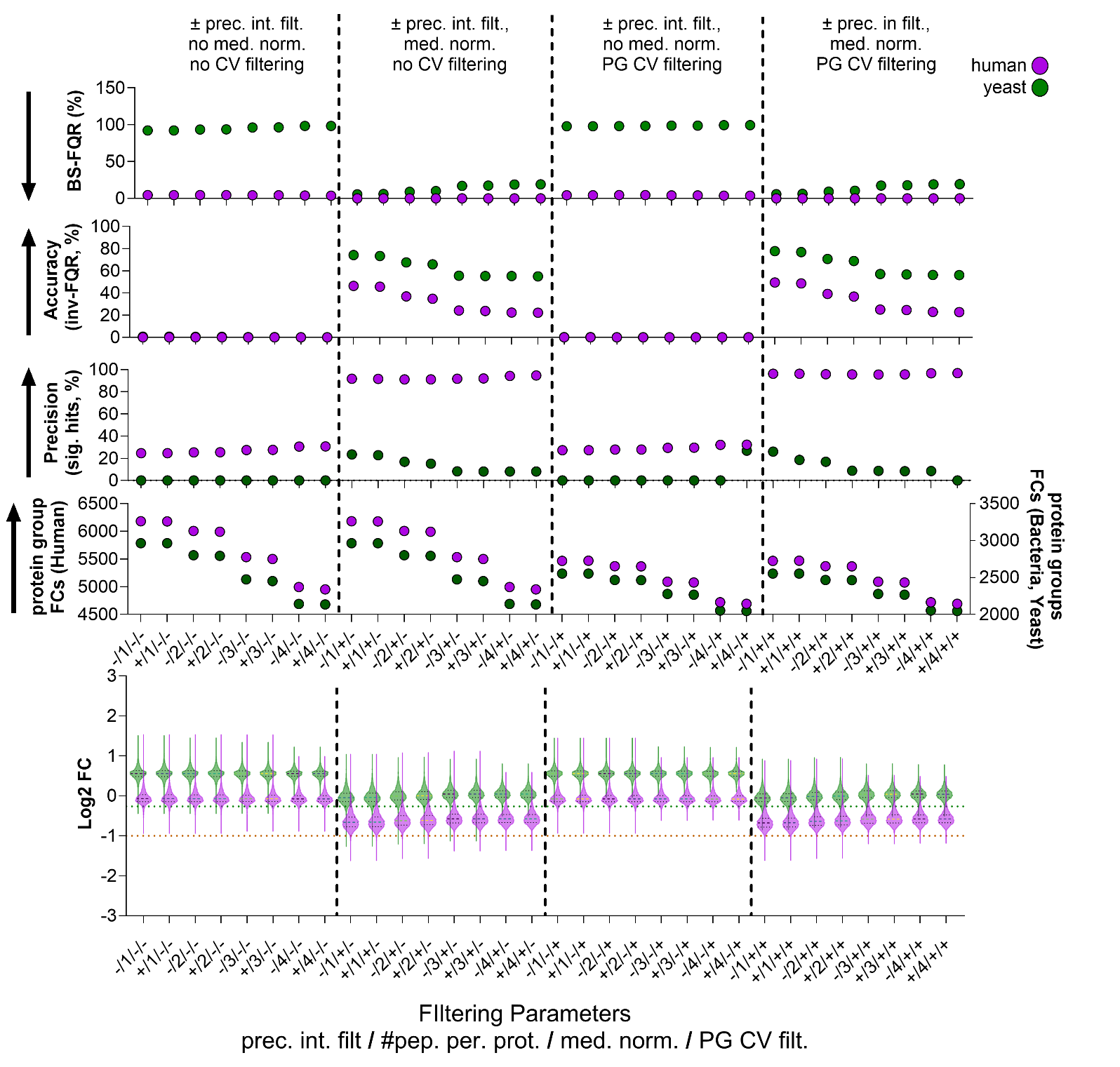

**C2**

**C3**

**C4**

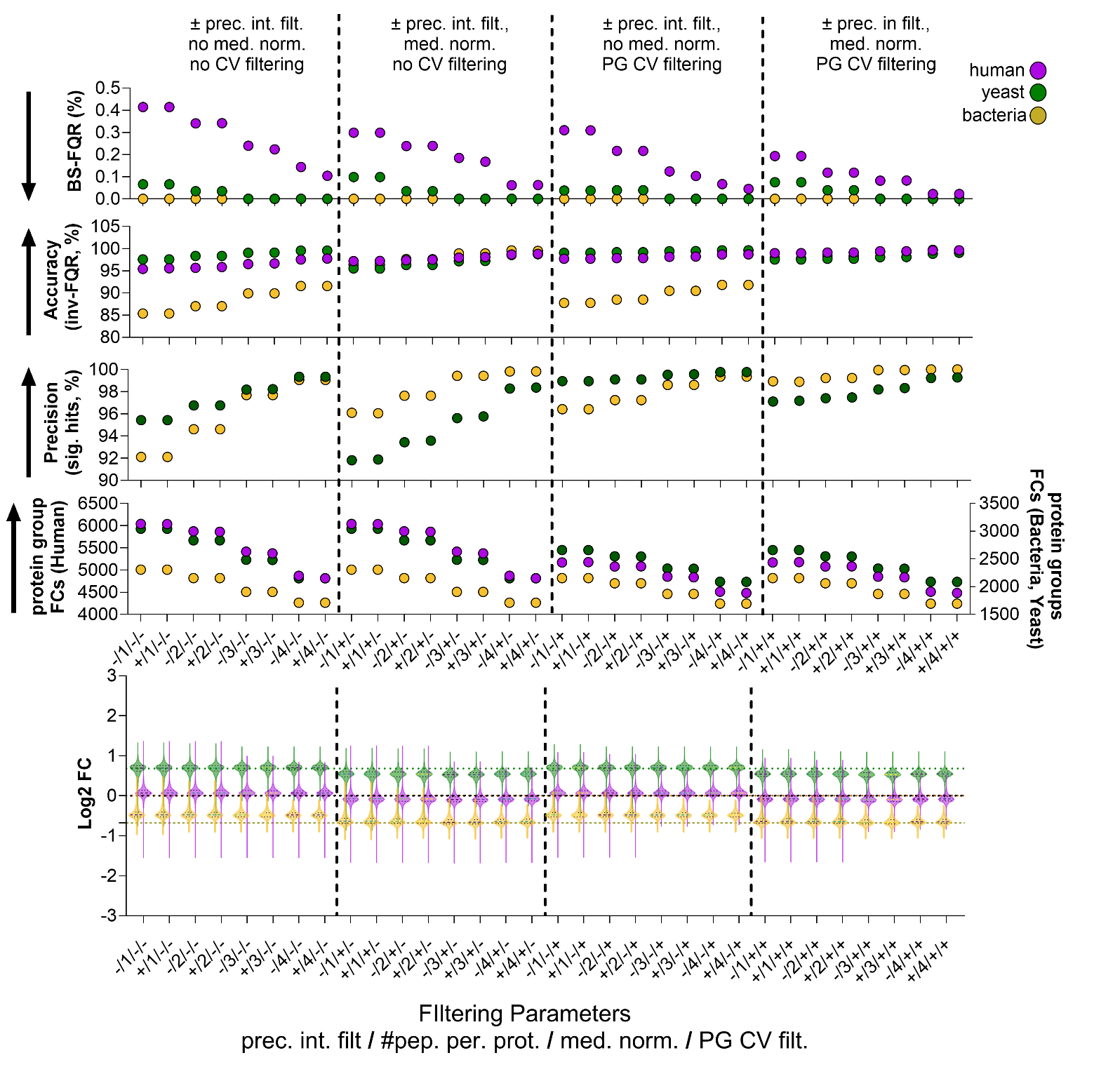

**C5**

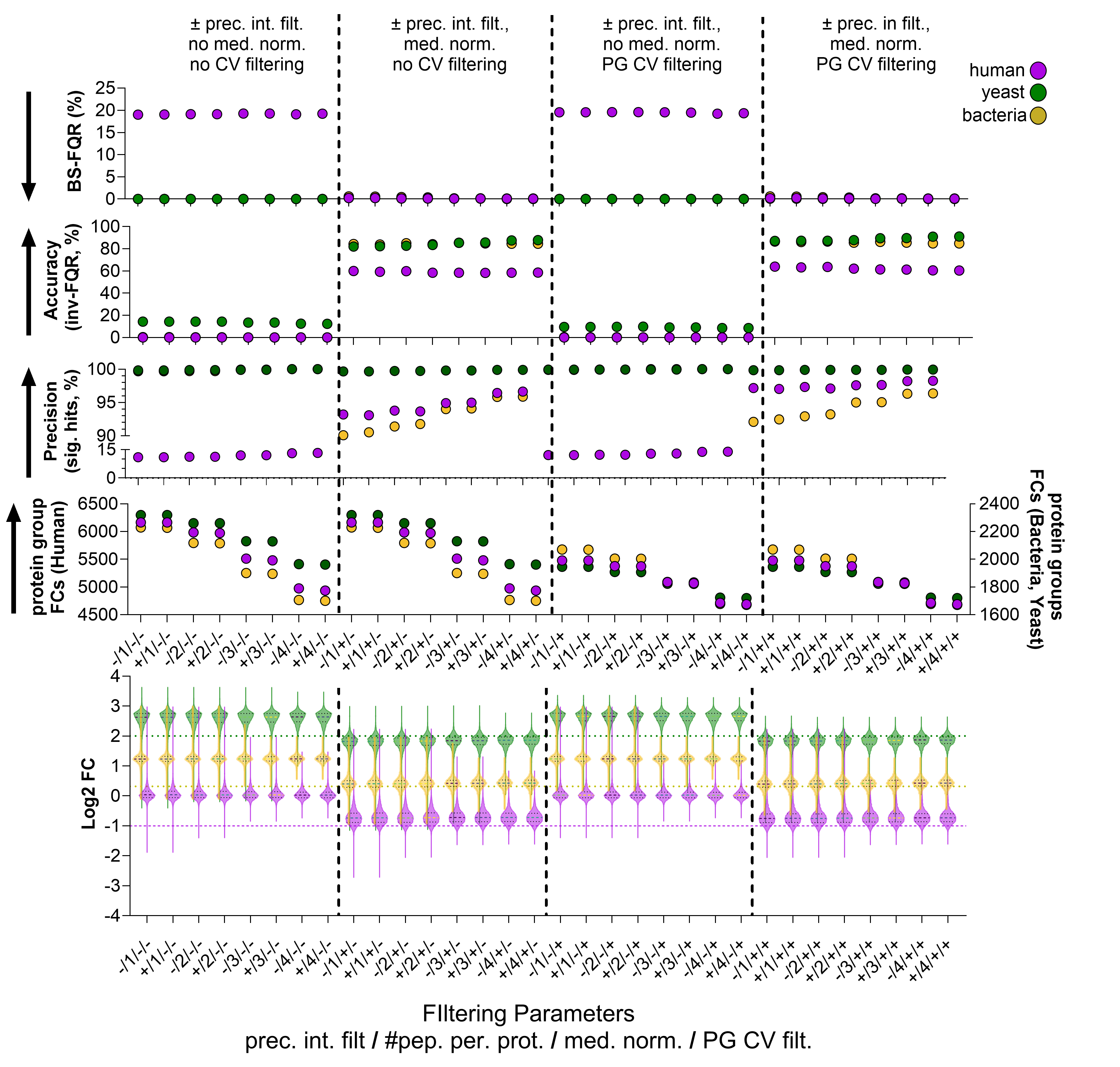

**C6**

**Supplemental figure 4.** **Post processing impacts quantitative efficacy in typical and atypical proteome comparisons derived from timsToF-HT analyses.** Purple = Human, Green= Yeast Yellow = Bacteria. Data shown are derived from five replicates. The comparisons (C1-C6) are indicated. Arrows indicate the preferred vector for each metric. Horizontal lines indicate the expected FC in the Log2FC distribution plot.

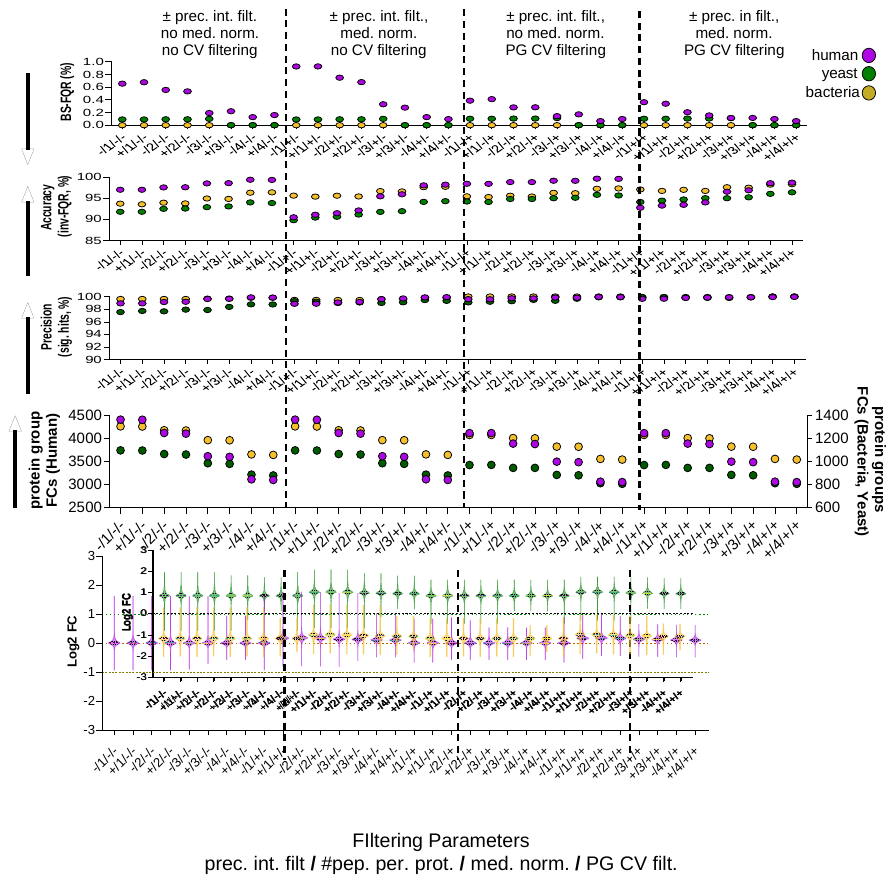

**C1**

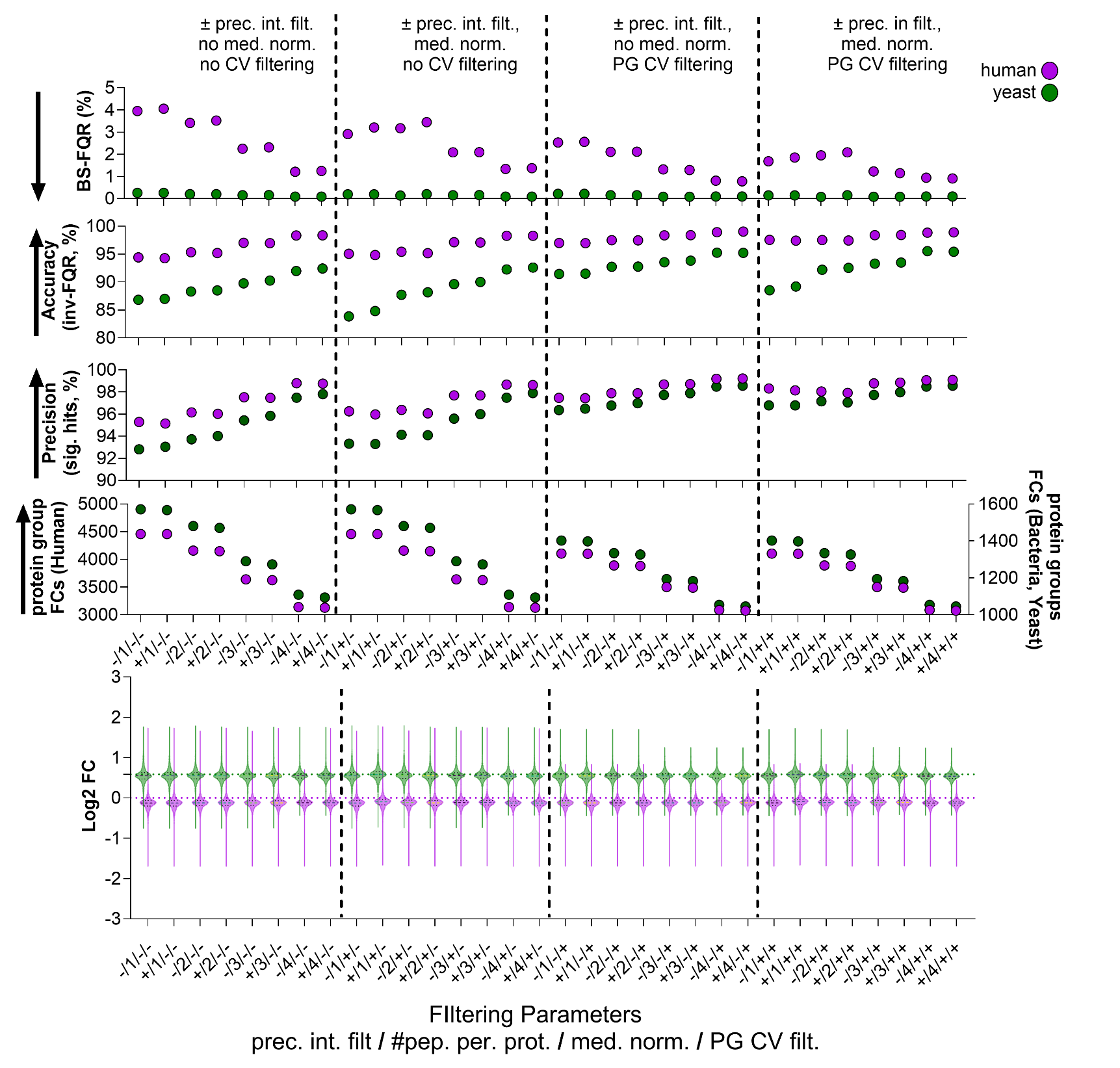

**C2**

**C3**

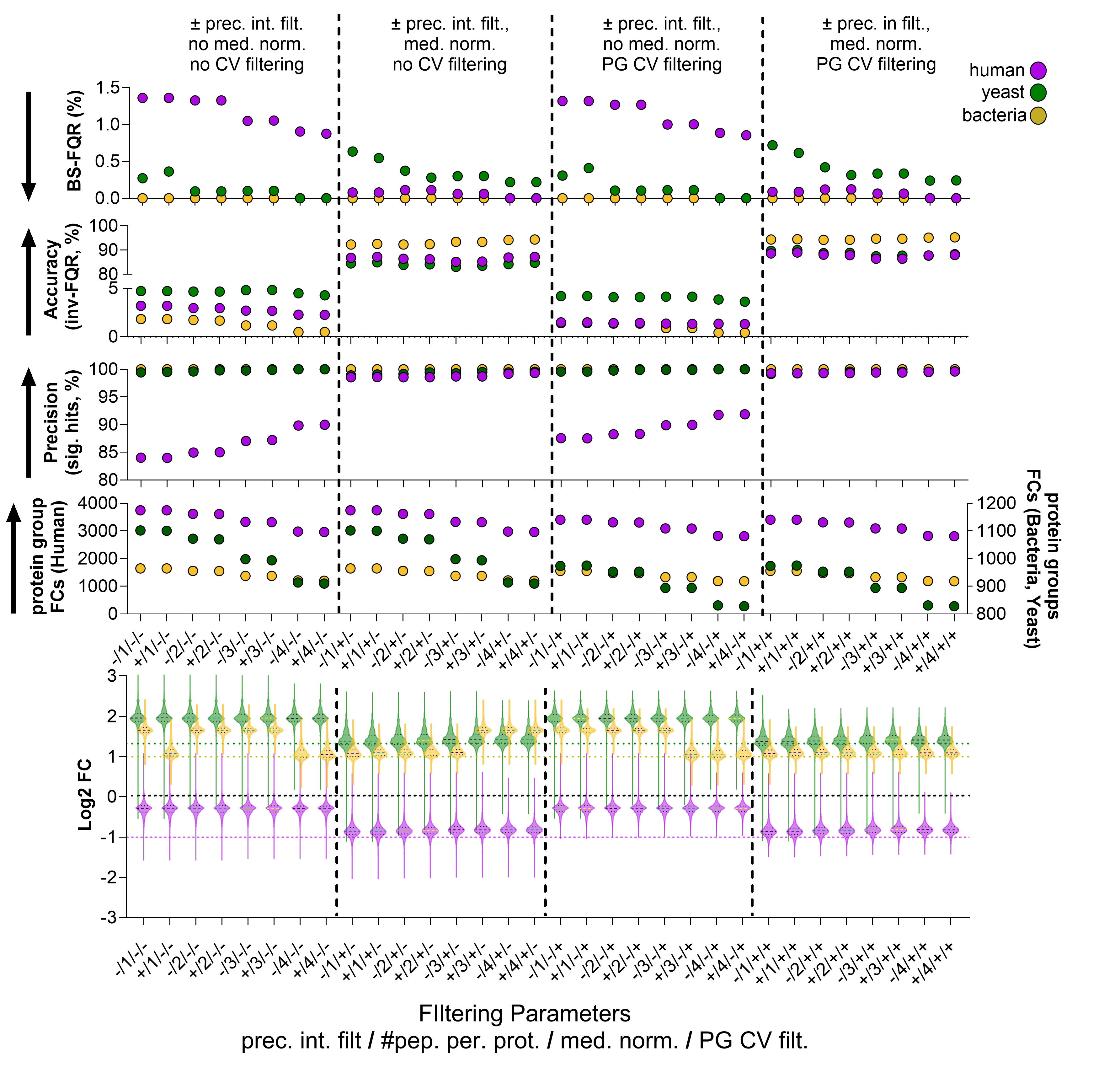

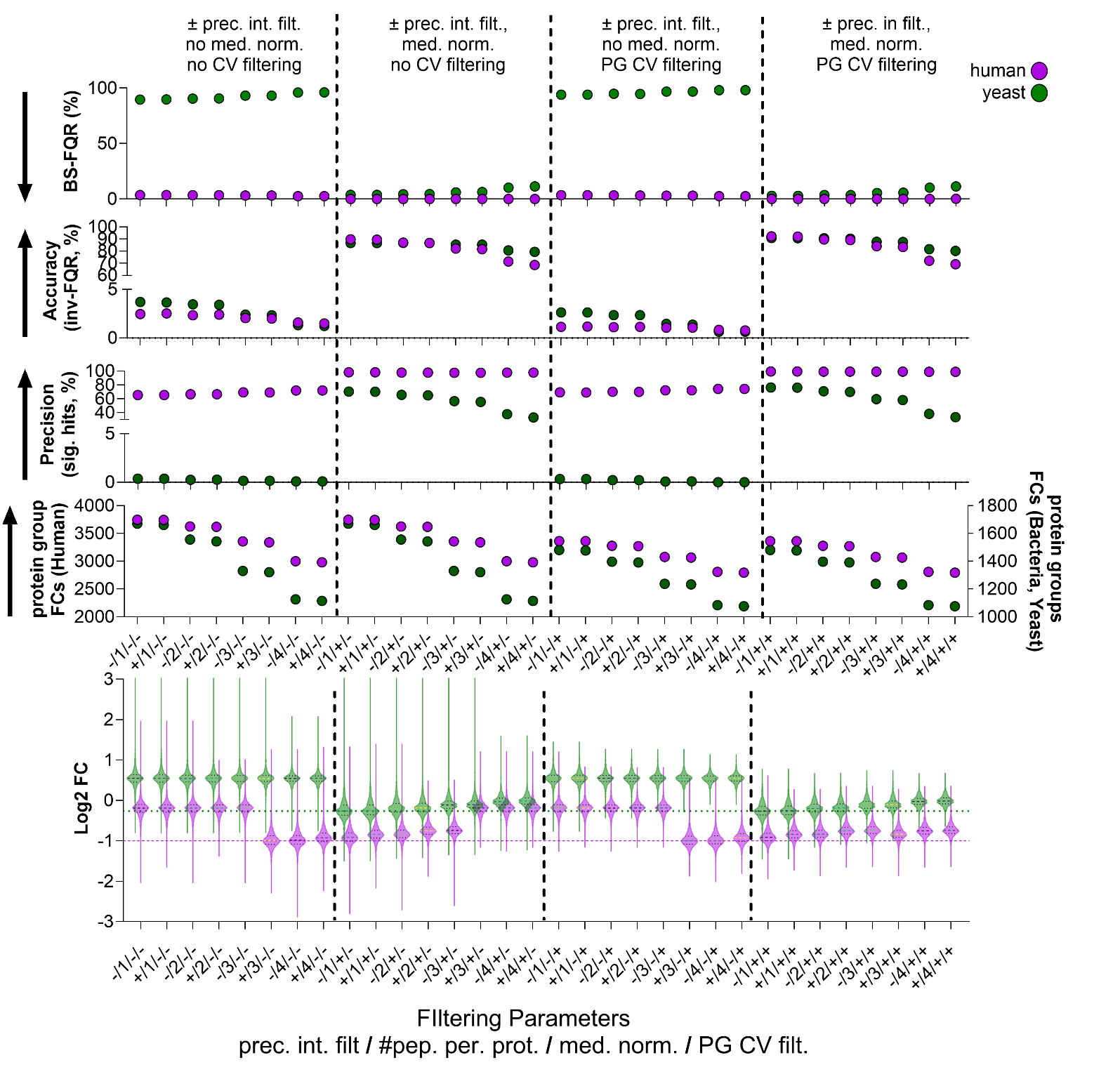

**C4**

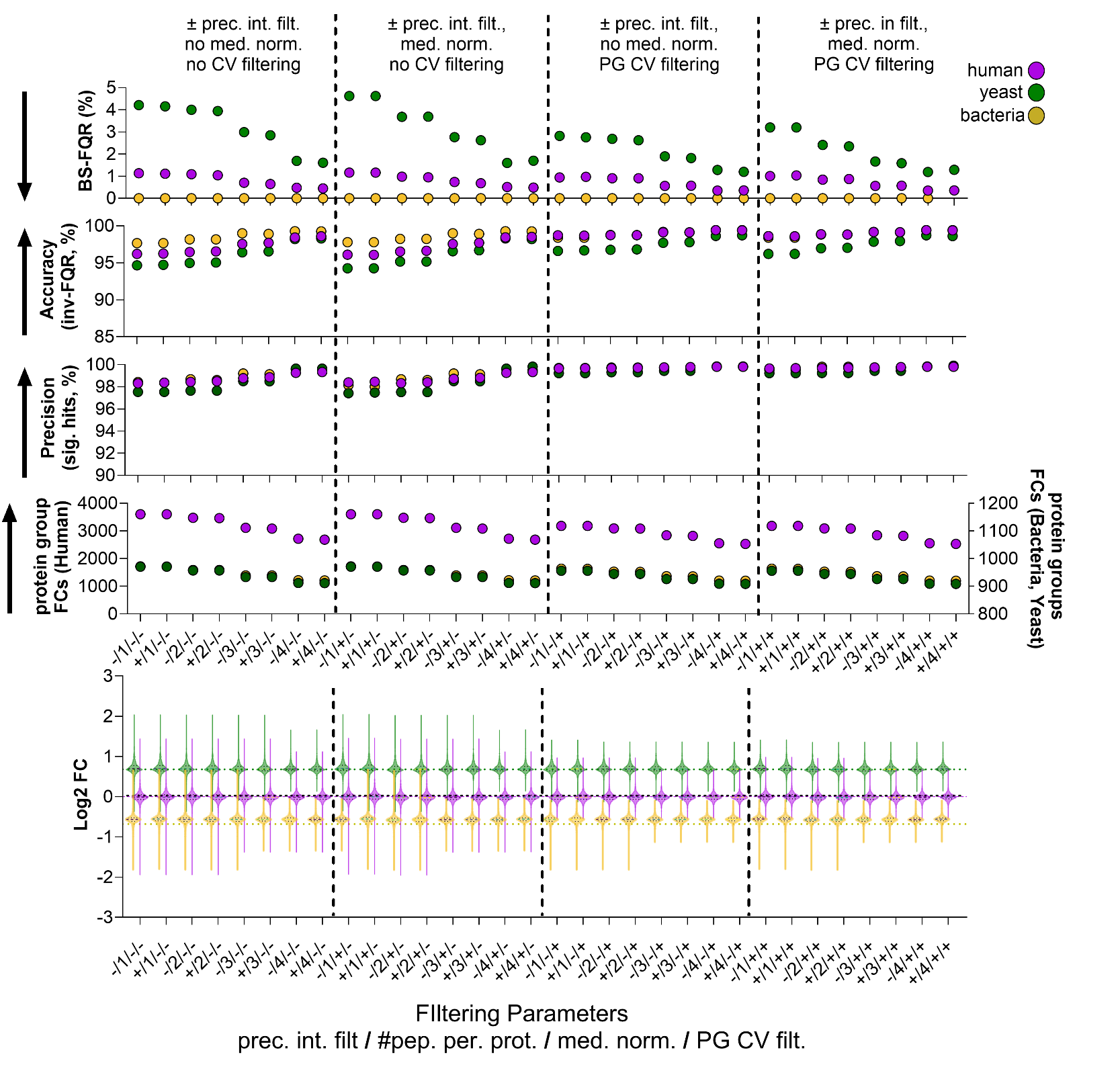

**C5**

**C6**

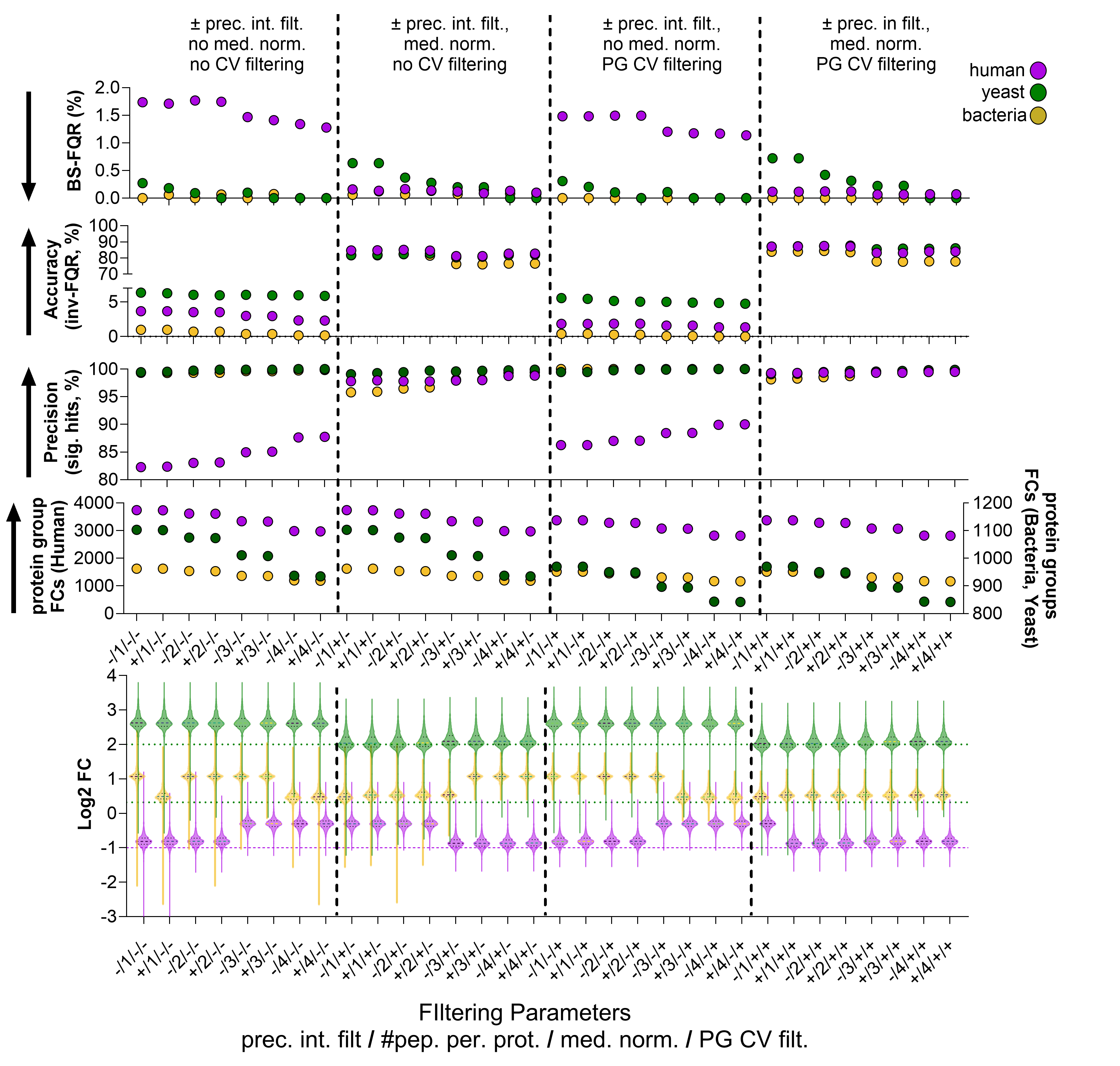

**Supplemental figure 5. Post processing impacts quantitative efficacy in typical and atypical proteome comparisons derived from Orbitrap Exploris 480 analyses.** Purple = Human, Green= Yeast Yellow = Bacteria. Data shown are derived from five replicates. The comparisons (C1-C6) are indicated. Arrows indicate the preferred vector for each metric. Horizontal lines indicate the expected FC in the Log2FC distribution plot.

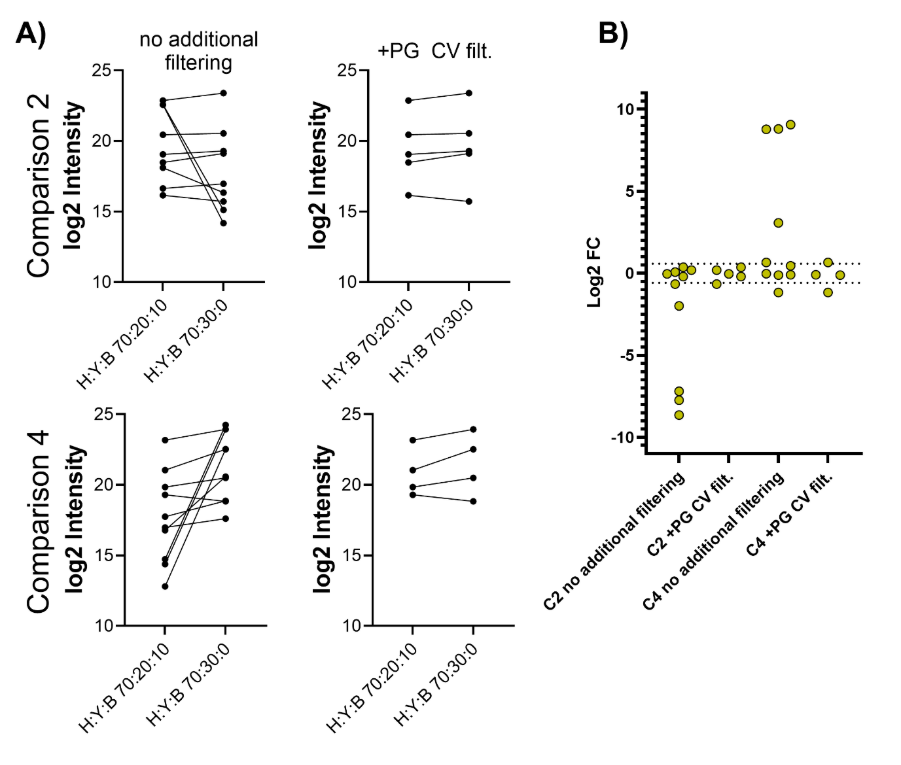

**Supplemental figure 6. Falsely-discovered changes in relative abundance are minimal, but may be present in typical proteomics analyses.** A) Intensity of the individual *E. coli* proteins quantified in comparisons 2 and 4, where one condition lacks *E. coli*. B) Log2 FC of *E. coli* proteins. Dotted lines indicate ±1.5-fold change. Data undergo either minimal post processing, or +protein group CV filtering as indicated. Data are derived from 5 replicates.

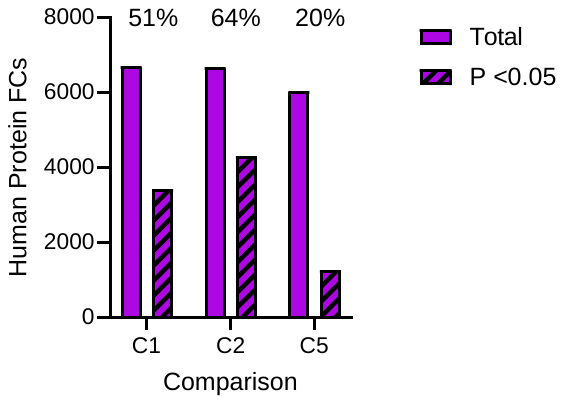

**Supplemental figure 7. Human proteins and those with statistical significance in comparisons with no ground-truth abundance change.** The number of human proteins relatively quantified in typical comparisons (C1, C2, C5) from timsToF experiments with minimal filtering are indicated, along with the number of these determined to be statistically significant (T-test with Benjamini Hochberg correction in Limma, adjusted P=<0.05). Percentages indicate the proportion of statistically significant proteins.

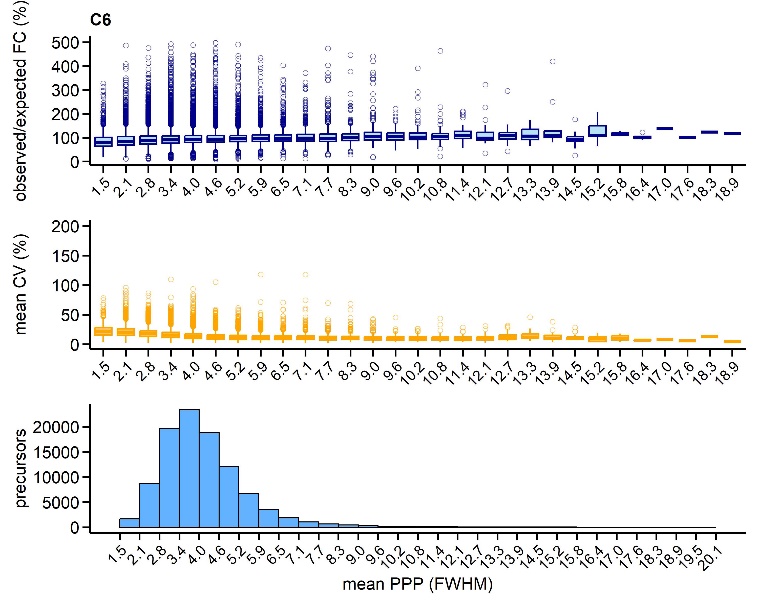

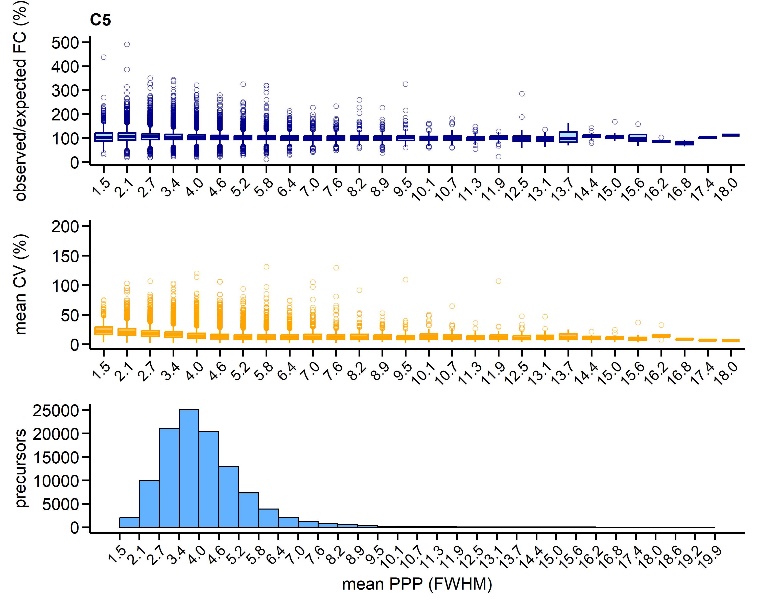

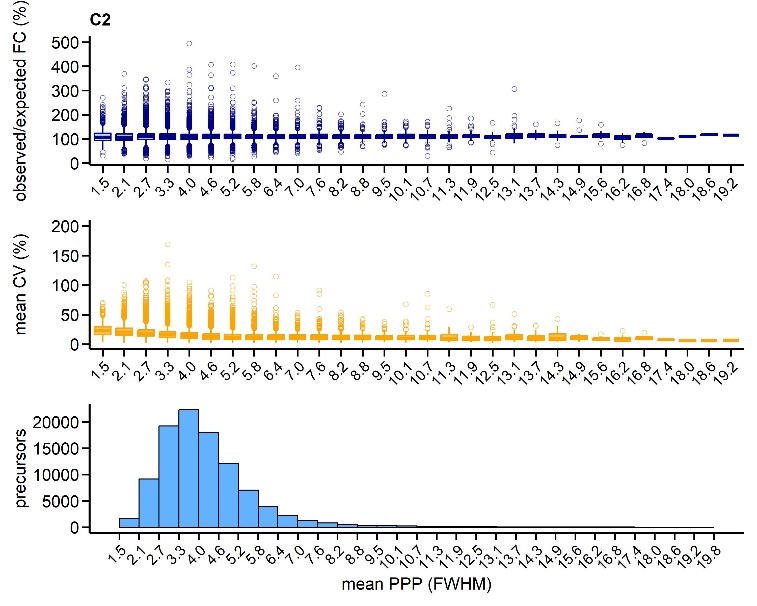

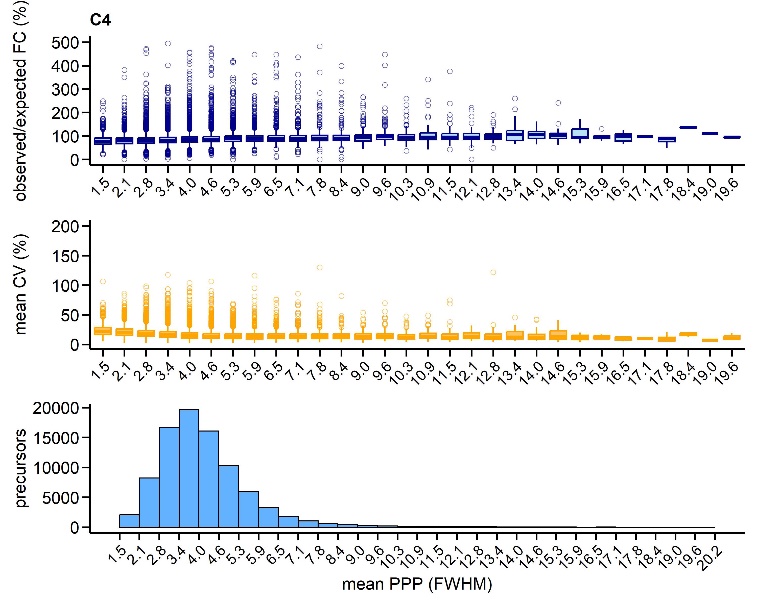

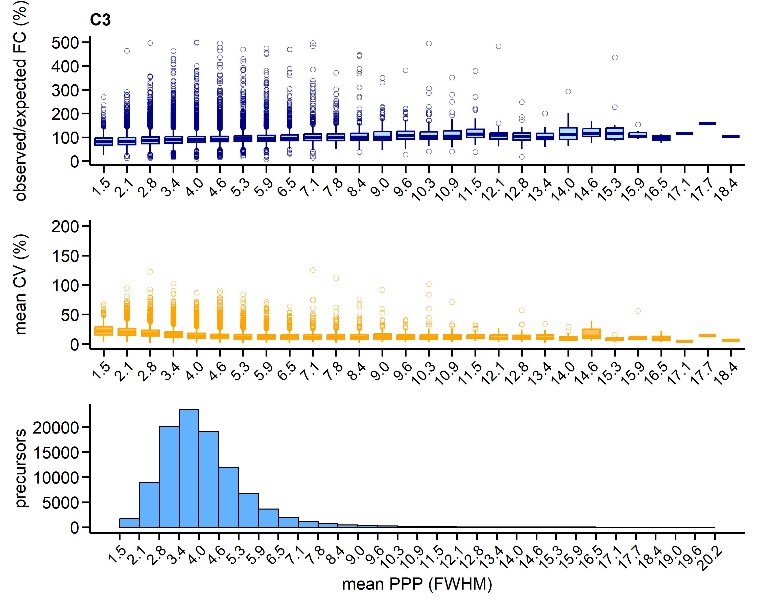

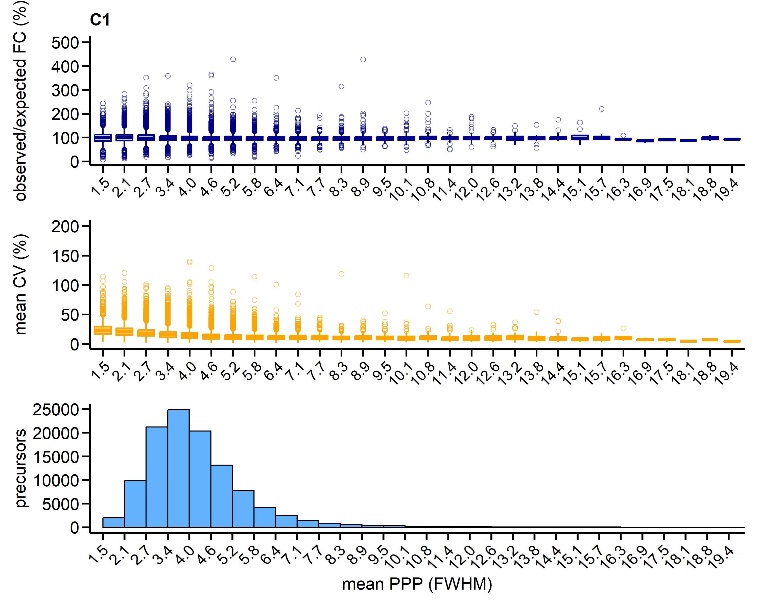
 **Supplemental figure 8. Relationship between precursor PPP (FWHM), precursor CV, and precursor deviation from ground truth.** timsToF precursors (with no filtering) were placed into 30 bins based on their PPP, for each comparison. For each precursor bin, the number of precursors was plotted-lowest histograms, as was the mean CV (%) and deviation from ground truth (observed/expected fold change, %)-middle and upper boxplots. Boxplots show median, quartiles, outliers. Data derived from 5 replicates.

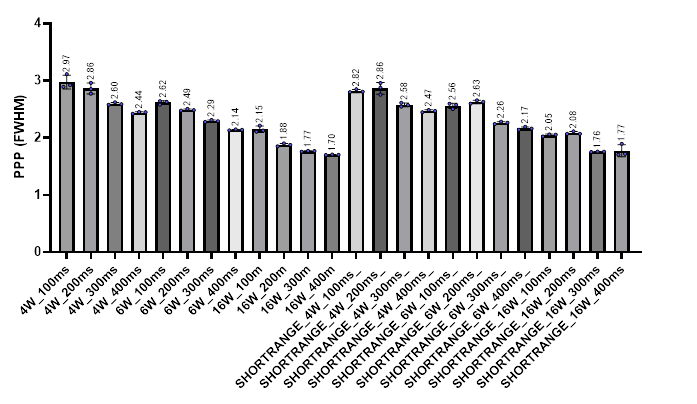

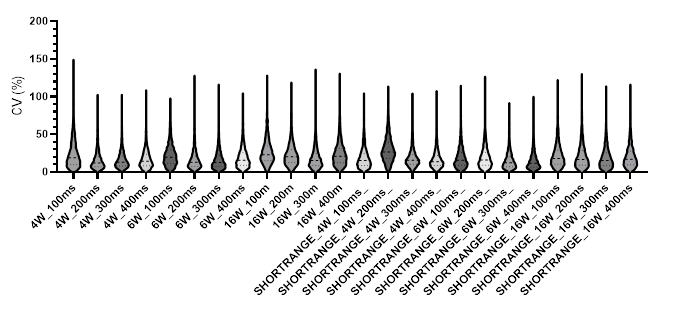

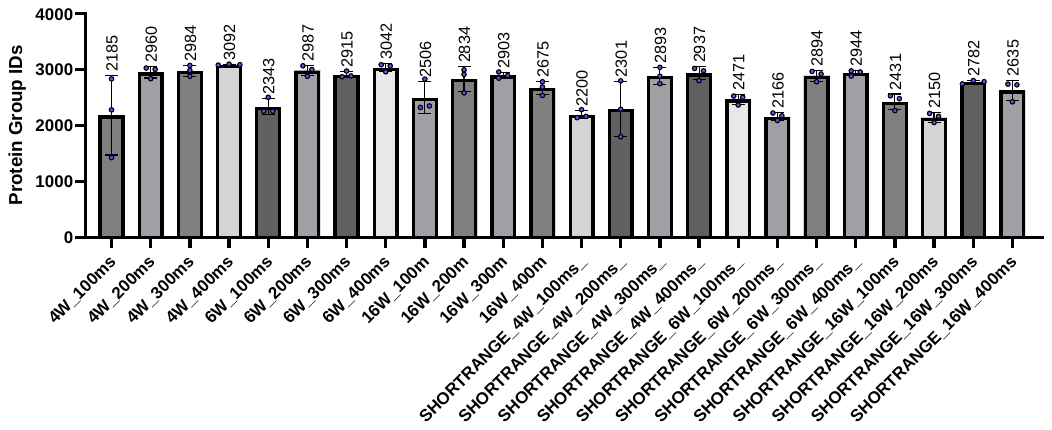

**Supplemental figure 9. High sensitivity diaPASEF optimisation-ID rates and reproducibility for single-cell level loads on the timsToF HT.** Tryptic HeLa digest-250pg-was subjected to different TARTs, PASEF window numbers, and mass ranges (*m/z* 300-1200 was considered default and not labeled, *m/z* 450-950 was short range). A) Protein group IDs, mean (bars) and replicate values (circles) shown. Error bars = SD. B) CV violin plot, dashed lines indicate median, and upper and lower quartiles. C) Points per peak (full width at half maximum), mean (bars) and replicate values (circles) shown. Error bars = SD.

**B)**

**C)**

**A)**
